# Metabolic response to point mutations reveals principles of modulation of *in vivo* enzyme activity and phenotype

**DOI:** 10.1101/738823

**Authors:** Sanchari Bhattacharyya, Shimon Bershtein, Bharat V. Adkar, Jaie Woodard, Eugene I. Shakhnovich

**Author notes:** Correspondence should be addressed to E.I.S. and S.Bh.

## Abstract

The relationship between sequence variation and phenotype is poorly understood. Here we use metabolomic analysis to elucidate the molecular mechanism underlying the filamentous phenotype of *E. coli* strains that carry destabilizing mutations in Dihydrofolate Reductase (DHFR). We find that partial loss of DHFR activity causes reversible filamentation despite SOS response indicative of DNA damage, in contrast to thymineless death (TLD) achieved by complete inhibition of DHFR activity by high concentrations of antibiotic Trimethoprim. This phenotype is triggered by a disproportionate drop in intracellular dTTP, which could not be explained by drop in dTMP based on the Michaelis-Menten like *in vitro* activity curve of Thymidylate Kinase (Tmk), a downstream enzyme that phosphorylates dTMP to dTDP. Instead, we show that a highly cooperative (Hill coefficient 2.5) *in vivo* activity of Tmk is the cause of suboptimal dTTP levels. dTMP supplementation rescues filamentation and restores *in vivo* Tmk kinetics to Michaelis-Menten. Overall, this study highlights the important role of cellular environment in sculpting enzymatic kinetics with system level implications for bacterial phenotype.

## Introduction

Understanding genotype-phenotype relationship is a central problem in modern biology. Mutations affect various layers of cellular organization, the mechanistic details of which remain far from being understood. Mutational effects propagate up the ladder of cellular organization from physico-chemical properties of biomolecules, up to cellular properties, finally affecting systems level properties, like the epigenome, transcriptome, proteome, metabolome or the microbiome. Collectively all layers in this multi-scale genotype-phenotype relationship dictates the fitness/phenotypic outcome of the mutations at the organism level. It has been shown, using the concept of a biophysical fitness landscape, that it is possible to predict fitness effects of mutations from a knowledge of molecular and cellular properties of biomolecules (Adkar *et al*, 2019; Adkar *et al*, 2017; Bershtein *et al*, 2015b; Dean & Thornton, 2007; Dykhuizen *et al*, 1987; Lunzer *et al*, 2005; Rodrigues *et al*, 2016) as well as using systems level properties like proteomics and transcriptomics (Bershtein *et al*, 2015a; Bershtein *et al*, 2017). The metabolome which is represented by the metabolite profile of the cell, is a more recent advancement in the -omics technology (Bennett *et al*, 2009). Metabolites represent end products of biochemical pathways; hence they are downstream to other -omics data, and therefore closest to the phenotype. Hence metabolomics is widely recognized now as an important stepping-stone to relate genotype to phenotype (Bhattacharyya *et al*, 2016; Bhattacharyya *et al*, 2018; Fiehn, 2002; Handakumbura *et al*, 2019; Harrison *et al*, 2020; Johnson *et al*, 2016; Patti *et al*, 2012; Rodrigues & Shakhnovich, 2019; Zampieri & Sauer, 2017). In the recent past, high-throughput studies have been dedicated to understanding how genetic variations lead to changes in metabolic profile of the cell (Fuhrer *et al*, 2017; Mulleder *et al*, 2016). Though vast knowledge is available in terms of how mutations perturb metabolite levels either in the local vicinity or distant in the network, a mechanistic knowledge of how such changes modulate phenotypic outcomes is lacking.

In this work, we use targeted metabolomics to understand the mechanistic basis of how destabilizing mutations in the essential core metabolic enzyme of *E. coli* Dihydrofolate Reductase cause pronounced (>10 times of the normal cells) filamentation of bacteria. DHFR catalyzes conversion of dihydrofolate to tetrahydrofolate, which is an essential one-carbon donor in purine, pyrimidine and amino acid biosynthesis pathways. Though filamentation is a documented effect of Trimethoprim (Tmp) (antibiotic that inhibits bacterial DHFR) treatment, it is mostly associated with thymineless death (TLD) (Sangurdekar *et al*, 2011) due to loss of thymidine nucleotides (dTTP) (Kwon *et al*, 2010) and consequent SOS response (Sangurdekar *et al*, 2010; Sangurdekar *et al*., 2011). In this study, although mutant DHFR strains incur a sharp drop in thymidine nucleotides (dTMP, dTDP and dTTP) and elicit SOS response, contrary to expectations, we found that filamentation is completely *reversible*. Using a range of Tmp concentrations, we delineated the conditions required for TLD which is complete inhibition of DHFR activity achieved by lethal doses of Tmp, while mutant DHFR strains only incur *partial* loss of activity. Additionally, we found that mutant strains have disproportionately low levels of dTDP (and hence dTTP) primarily due to the strongly cooperative *in vivo* activity of the downstream essential pyrimidine biosynthesis enzyme Thymidylate Kinase (Tmk), which phosphorylates dTMP to dTDP. This is in stark contrast to its Michaelis-Menten (MM) activity profile observed *in vitro* and is therefore manifested in terms of reduced recovery of intracellular dTDP/dTTP even as dTMP begins to increase. Surprisingly, supplementation of external dTMP in the medium which rescues *in vivo* dTDP levels and filamentation, switches the *in vivo* Tmk activity curves to the conventional ‘*in vitro* like’ MM kinetics. The cooperative enzyme activity is best explained by diffusion-limitation of substrate dTMP *in vivo*, possibly due to substrate channeling and metabolon formation. This idea is also supported by Hill-like and hyperbolic *in vivo* activity curves of a number of other *E. coli* proteins. Overall, this study highlights the pleiotropic nature of mutations and the way in which the complex cellular environment and metabolic network modulates *in vivo* enzyme activity and organismal fitness.

## Results

### Several chromosomal mutations in folA gene give rise to slow growth and filamentation of E. coli

Earlier we had designed a group of highly destabilizing chromosomal DHFR mutations in *E. coli* MG1655 (W133V, V75H+I155A, I91L+W133V, and V75H+I91L+I155A) that cause very slow growth at 37°C and 42°C ((Bershtein *et al*, 2013; Bershtein *et al*, 2012) and Figure 1A, see Methods for details about the strains). While WT, W133V and V75H+I155A could grow up to 42°C, mutants I91L+W133V and V75H+I91L+I155A could only survive up to 40°C. To understand the effects of these mutations on bacterial morphology, we grew mutant DHFR strains under two different growth conditions: M9 minimal media without and with supplementation with casamino acids (mixtures of all amino acids, except tryptophan, see *Methods*). In minimal media, median cell lengths of some mutants (W133V and V75H+I155A at 42°C) were smaller compared to WT, while I91L+W133V (at 40°C) was marginally longer than WT (Figure 1C). However, when M9 minimal medium was supplemented with amino acids, we found that cells carrying these mutations were pronouncedly filamentous. Figure 1B shows live cell DIC images of wild-type (WT) and I91L+W133V mutant DHFR strains at 30°C, 37°C, and 42°C (40°C for I91L+W133V strain). (See Supplementary Figure EV1A for images of other low fitness mutant strains). In parallel to the detrimental effect of temperature on fitness, we noted that the morphologies were also temperature sensitive. I91L+W133V and V75H+I91L+I155A strains exhibited a 1.5-1.75 fold increase (comparatively to WT) in the average cell length at 37°C (Figure 1D), while W133V and V75H+I155A were not elongated at 37°C. The latter, however, showed an increase up to 2.0-2.3 fold over WT cell lengths at 42°C (Figure 1E). Strains I91L+W133V and V75H+I91L+I155A showed 1.8-2.0 fold increase in the average cell length at 40°C, with some cells reaching up to 20μm in length (about 10 fold increase). Besides temperature of growth, since filamentation was also strongly dependent on availability of amino acids in the growth medium, it seemed likely that it was the result of a metabolic response due to partial loss of DHFR function.

**Figure 1:**
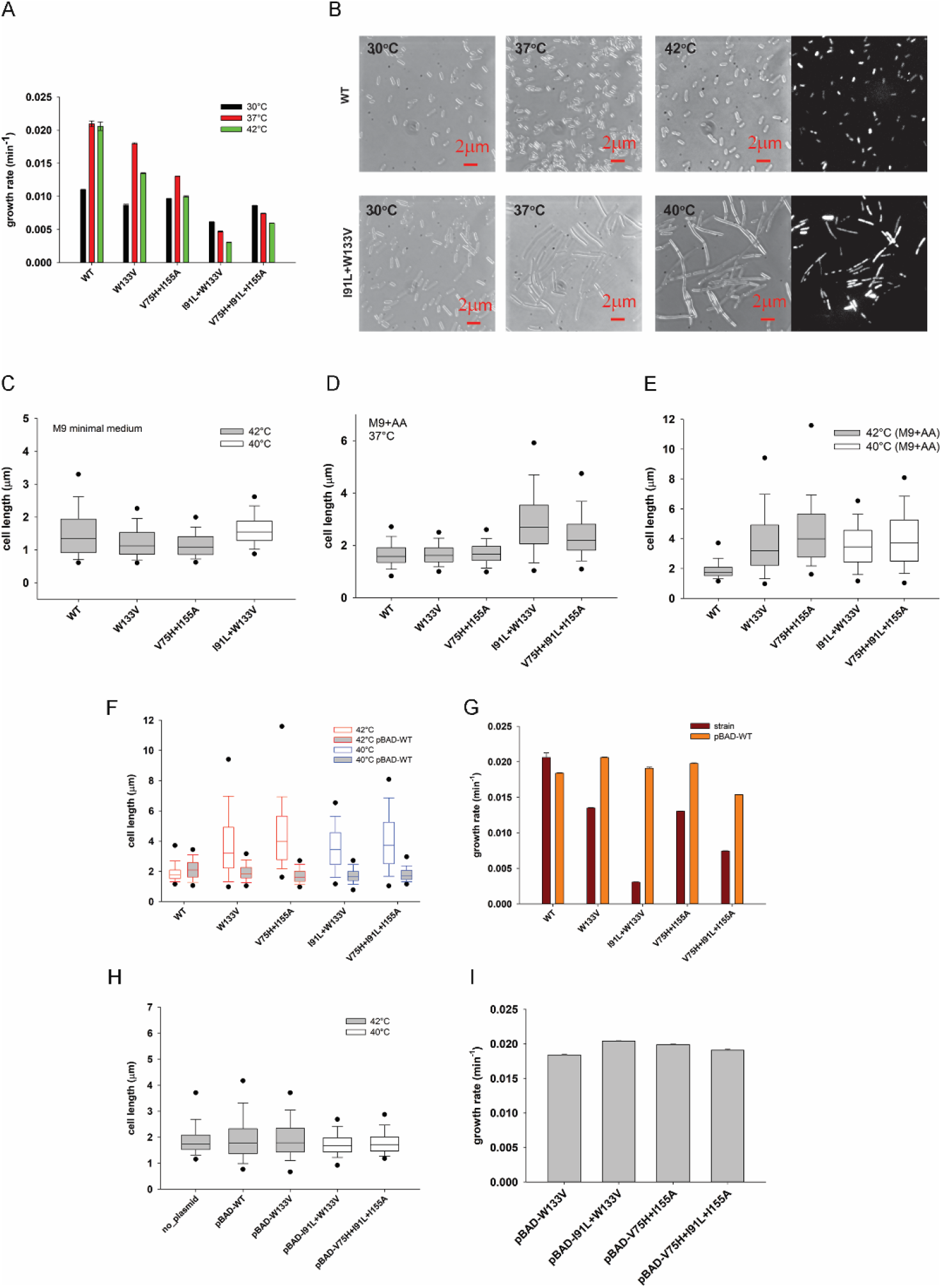
Destabilizing mutations in DHFR induce filamentous phenotype due to loss of DHFR activity. (A) Growth rates of mutant DHFR strains at 30°C, 37°C and 42°C. While most mutants grow well at 30°C, they grow very poorly at high temperatures. Error bars represent SEM of three biological replicates. (B) Live cells DIC images and DAPI nucleoids staining of WT DHFR and I91L+W133V DHFR strains after being grown at 30°C, 37°C, or 42°C (I91L+W133V was grown at 40°C) in M9 medium supplemented with amino acids for 4 hours (see Methods). Cell lengths were measured from the obtained DIC images (see Methods) and their distribution at 37°C and 40°C/42°C is shown in (C) and (D) as box-plots (see Methods). Images of other mutant DHFR strains W133V, V75H+I155A and V75H+I91L+I155A are presented in related Figure EV1A. (E) Distribution of cell lengths of WT and mutants W133V and V75H+I155A at 42°C (gray box) and I91L+W133V at 40°C (represented as white box) after being grown in M9 minimal medium (without amino acids) for 4 hours. Median cell length of W133V and V75H+I155A is significantly smaller than WT (Mann-Whitney test, p-value <0.001). Functional complementation of WT DHFR from an arabinose inducible pBAD plasmid rescues both (F) filamentation and (G) growth defects of mutant strains grown at 42°C (for WT, W133V and V75H+I155A strains) or at 40°C (for I91L+W133V and V75H+I91L+I155A) in M9 medium supplemented with amino acids. Expression of mutant proteins from pBAD plasmid on the WT background does not result in (H) filamentation or (I) growth defects. For the boxplots in panels C, D, E, F and H, the central band represents the median of the distribution, the box ends represent the 25^th^ and 75^th^ percentile, the whiskers represent the 10^th^ and 90^th^ percentile, while the dots represent the 5^th^ and 95^th^ percentile. Data was usually obtained from 2-3 biological replicates. The number of cells used to derive the boxplot distributions in the different panels range usually between 200 to 450 (please refer to Figure 1 source data for exact number of cells for each dataset). See related Figure EV1B.

### Filamentation is due to drop in DHFR activity

DHFR is a central metabolic enzyme that is involved in conversion of dihydrofolate to tetrahydrofolate, and the latter is an important 1-carbon donor in the biosynthesis of purines, pyrimidines, and certain amino acids like glycine and methionine. Earlier we had reported that these mutant DHFR strains had very low abundance of the mutant proteins in the cell ((Bershtein *et al*., 2015a; Bershtein *et al*., 2013) and Figure EV1B), an effect that could be rescued by deletion of Lon protease or by over-expressing chaperones like GroEL-ES (Bershtein *et al*., 2013). We, therefore, reasoned that filamentation could be a result of drop in DHFR activity in these cells. To confirm this, we supplemented the *E. coli* strains carrying chromosomal DHFR mutations with WT DHFR expressed from a plasmid and found that both filamentation (Figure 1F) and growth defects (Figure 1G) were fully rescued. On the WT background, expression of extra DHFR resulted in some elongation, presumably due to toxicity of DHFR over-expression (Bhattacharyya *et al*., 2016). We also found that plasmid expression of mutant proteins in WT cells did not result in any filamentation (Figure 1H) or growth defects (Figure 1I). This shows that filamentation is not due to toxicity of the mutant DHFR proteins. We also found that treatment of WT cells with Trimethoprim, an antibiotic that targets bacterial DHFR (Figure EV1C), also caused filamentation at concentrations near the MIC (1μg/ml). At higher concentrations of the drug, there is growth arrest, hence no filamentation, leading to a non-monotonic dependence of cell length on Tmp concentration (Figure EV1, panels D and E at 37°C and 42°C respectively).

### Filamentous strains exhibit imbalance between dTTP and other deoxyribonucleotides

Here we aimed to determine metabolic changes in mutant strains associated with filamentous phenotype. To that end we carried out metabolomics analysis under conditions of filamentation (in amino acid supplemented M9 medium) as well as under non-filamentation conditions (in minimal medium). As shown in Figure 1, W133V and V75H+I155A mutants behave similarly (in terms of growth rates, extent of filamentation and ability to grow up to 42°C), while mutants I91L+W133V and V75H+I91L+I155A behaved similarly. Moreover, I91L+W133V had the lowest growth rate among all mutants, which makes it an ideal candidate to study the extreme effect of mutations. Hence, for metabolomics analyses as well as for several later experiments, we chose one representative example from each of these two groups, namely W133V, I91L+W133V, as well as WT treated with 0.5μg/ml Tmp.

We observed that in the absence of amino acids, *when the cells are not filamented* mutant strains as well as WT cells treated with 0.5μg/ml Tmp (close to MIC) exhibited very low levels of both purines and pyrimidines (Figure 2A). For example, in strain I91L+W133V, IMP, AMP and dTMP levels were respectively 17%, 30% and 5% of WT levels, while dTTP levels were below the detection limit. Methionine and glycine biosynthesis require tetrahydrofolate derivatives, hence, expectedly, methionine levels were only 1-3% in mutant strains (Figure 2B). Overall, we conclude that large drop in methionine and purine (IMP) levels presumably stalls protein/RNA synthesis. Since increase in cell mass is essential for filamentation, cells under this condition are not filamented.

**Figure 2:**
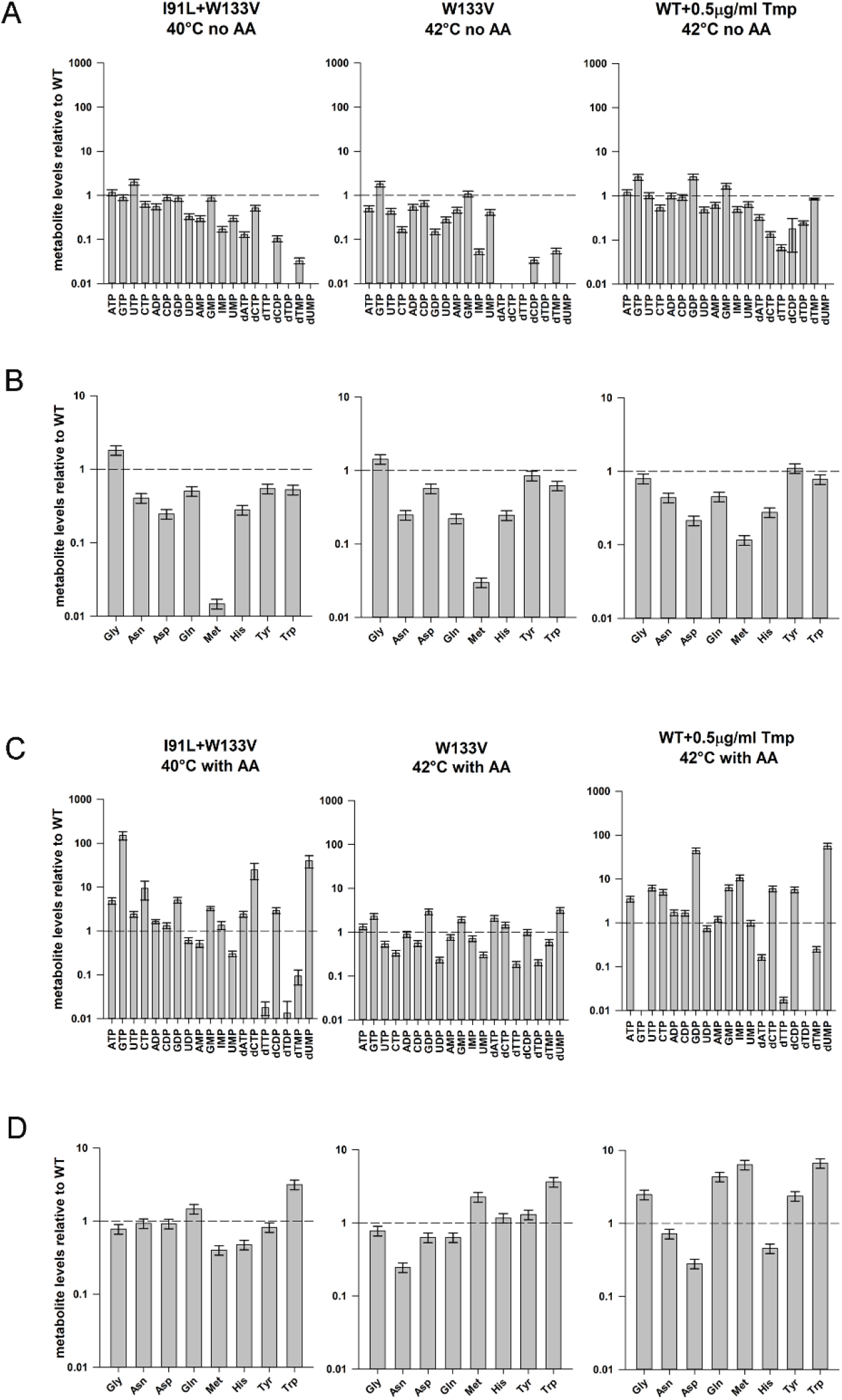
Metabolomics of mutant DHFR strains in minimal media without or with added amino acids. (A) and (B) shows abundance of selected nucleotides and amino acids for mutants I91L+W133V and W133V as well as WT strain treated with 0.5 μg/ml of Tmp (Trimethoprim) after 4 hours of growth at 40/42°C in M9 minimal medium (condition of no filamentation), while (C) and (D) represents nucleotide and amino acid abundances after 4 hours of growth in amino acid supplemented M9 medium at 40/42°C (condition of filamentation). Concentration of all metabolites were normalized to WT levels at 4 hours when grown under similar conditions. In minimal medium (B), Methionine levels are extremely low, which recover largely in panel (D). Levels of purines (IMP, AMP) are also largely rescued with amino acid supplementation, however dTMP, dTDP and dTTP levels remain extremely low. Error bars represent SEM of at least three biological replicates (see Methods). The dashed line in each plot represents value of 1 for WT.

In the presence of 1% casamino acids, several but not all amino acids showed a marked increase in abundance (Figure 2D). Methionine levels rose to 40% of WT levels for I91L+W133V mutant, while aspartate/asparagine, glutamine, histidine and tryptophan levels also showed a significant increase. Particularly interesting was the fact that purine levels were substantially rescued upon addition of amino acids (Figure 2C). IMP showed the maximal effect, increasing 10-15 fold over its levels in the absence of amino acids. ATP, ADP, AMP and GMP also showed similar trends. Since the product of DHFR is eventually used in the synthesis of methionine, IMP and dTMP (Figure EV2A), we hypothesize that addition of methionine in the medium allows higher amounts of 5,10-methylene-THF to be channeled towards synthesis of purines and pyrimidines. Moreover, both de novo purine and pyrimidine biosynthesis pathways require aspartate and glutamine (Figure EV2B, C), which were otherwise low in minimal medium. Overall, the metabolomics data suggest that in the presence of added amino acids, protein, and RNA synthesis are no longer stalled, and therefore growth, which is pre-requisite for filamentation, can happen.

Though dTMP levels increased to about ~10% of WT levels in I91L+W133V and ~50% in W133V, surprisingly, dTTP levels (thymidine derivative that is incorporated in the DNA) were only about 1% of WT levels in I91L+W133V and 18% for W133V (Figure 2C). In contrast, dATP and dCTP levels were very high (Figure 2C). We hypothesize that as cellular growth continues, misbalance in the concentrations of deoxy nucleotides may lead to erroneous DNA replication, induction of SOS response and blocked cell division. Indeed, in our previous study, proteomics and transcriptomics analyses showed that several SOS response genes were upregulated in I91L+W133V and V75H+I91L+I155A strains at 37°C (Bershtein *et al*., 2015a). We note that the imbalance of deoxy nucleotides due to extremely low dTTP levels happens even in the absence of added amino acids (Figure 2A), however this may not cause erroneous DNA replication as the cell cannot duplicate its cellular mass due to depletion of methionine and purines (so-called stringent response), which are essential for protein and RNA synthesis, and hence does not attempt cell division. Therefore this condition does not result in filamentation.

### Filamentation and SOS response: Deletion of recA rescues filamentation

We quantified the expression of several SOS genes *recA, recN* and *sulA* under different supplementation conditions for mutant DHFR strains as well as Tmp-treated WT cells. Mutants grown at low temperature and those grown in the absence of amino acids were not elongated, and not surprisingly, they did not elicit any SOS response (Figure 3A-C). In comparison, for all conditions that are associated with filamentation (I91L+W133V mutant at 37°C, W133V and I91L+W133V at 42°C) induced strong SOS response (Figure 3A-C). The levels of induction of these genes were greatly reduced in the presence of dTMP, consistent with lack of filamentation under this condition (Figure 3A-C). Similar trends were also observed for WT treated with Tmp

**Figure 3:**
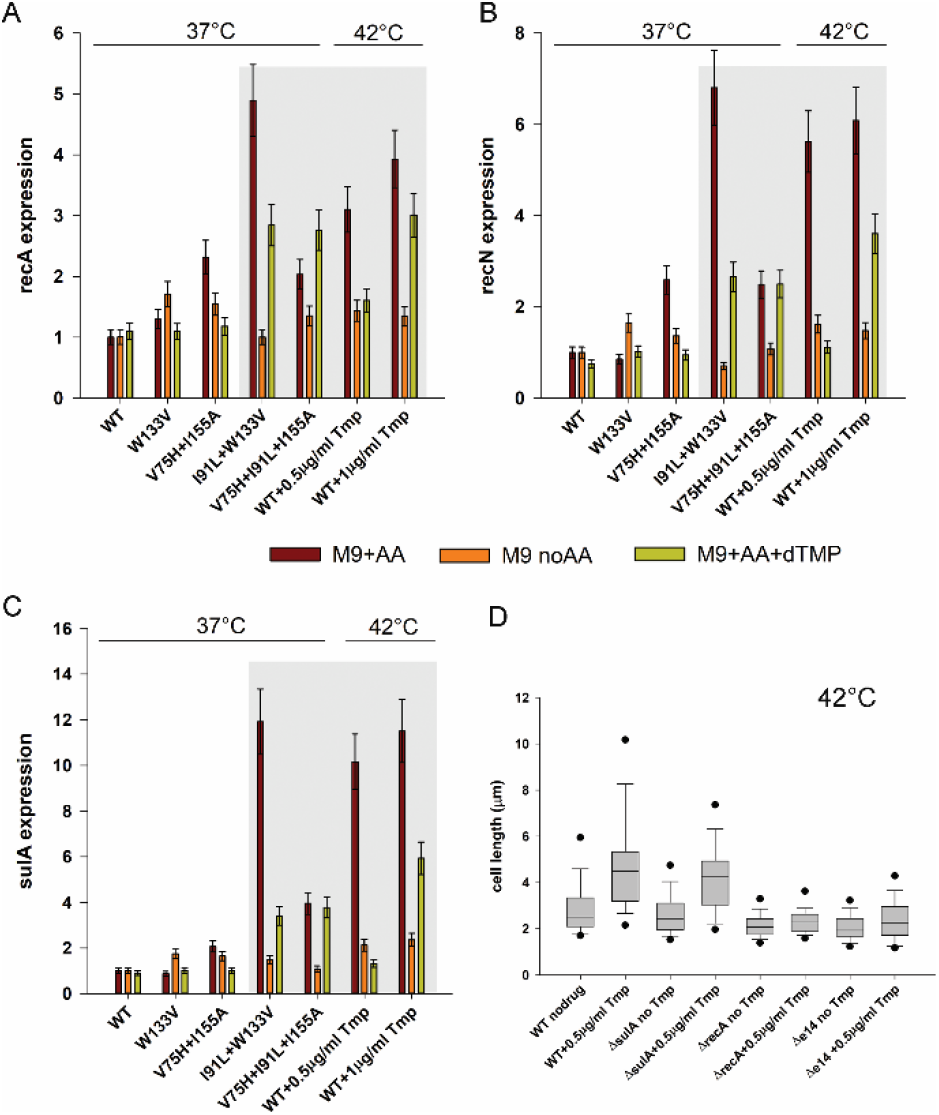
Filamentation in mutant DHFR strains is associated with strong SOS response. Expression of (A) recA (B) recN and (C) sulA genes measured by quantitative PCR when WT and mutant strains are grown in M9 medium with or without supplementation of amino acids or dTMP. WT and mutant strains were grown for 4 hours of growth in the indicated medium at 37°C, while WT treated with different concentrations of Tmp were grown for 4 hours at 42°C. Brown bars (M9+AA) in the gray shaded area correspond to filamentation conditions and these are associated with pronounced upregulation of all three SOS genes. On the other hand, conditions with loss of filamentation (with dTMP or no supplementation) show much less expression. Error bars represent SD of 2-3 biological replicates (see Methods). (D) Treatment of WT E. coli cells with sub-MIC concentration of Tmp (0.5 μg/ml) leads to filamentation at 42°C when grown in amino acid supplemented medium. However, a recA knock-out strain under similar condition shows no elongation, indicating the role of SOS pathway in filamentation. A sulA knock-out continues to elongate, indicating the role of sulA-independent pathways. An E. coli strain deleted for the e14 prophage region however showed no filamentation upon Tmp treatment, indicating that sfiC gene in the e14 region might be one such sulA independent player. The central band in the box plots represents the median of the distribution, the box ends represent the 25^th^ and 75^th^ percentile, the whiskers represent the 10^th^ and 90^th^ percentile, while the dots represent the 5^th^ and 95^th^ percentile. Data was usually obtained from 2-3 biological replicates. The number of cells used to derive the boxplot distributions in the different panels range usually between 100 to 200 (please refer to Figure 3 source data for exact number of cells for each dataset).

Overexpression of RecA is the key trigger of the SOS response to DNA damage. RecA cleaves the dimeric LexA repressor to turn on the genes that are under the SOS box (e.g., sulA, uvr proteins, *etc*.). SulA inhibits the cell division protein FtsZ, eventually causing filamentation. To understand the role of *recA* and *sulA* in our study, we treated *ΔrecA* and *ΔsulA* strains with near-MIC levels of the antibiotic Tmp. As expected, *ΔrecA* strain did not show filamentation upon Tmp treatment (Figure 3D), clearly highlighting its definitive role in filamentation. However, *ΔsulA* strain continued to filament (Figure 3D), suggesting possible role of sulA-independent pathways. This data, however, does not negate out the role of sulA, since sulA is highly upregulated in mutant strains and is a well-known inhibitor of the cell division protein FtsZ. A poorly characterized segment of the *E. coli* genome called the e14 prophage that harbors the *sfiC* gene has been implicated to play a role in SOS dependent but sulA independent filamentation (Maguin *et al*, 1986). To find out if this e14 prophage region and the sfiC gene are involved in a sulA independent SOS pathway in our case, we used a mutant *E. coli* strain that is knocked out for the e14 prophage region and found that this strain does not filament upon Tmp treatment (Figure 3D), indicating that *sfiC* might be one of the players involved in elongation, though the exact mechanism of action through this pathway is unclear. Regardless of the mechanism, these results clearly establish the role of the SOS pathway towards filamentation in mutant DHFR strains, which in turn is due to imbalance between dTTP and other deoxynucleotides in the cell.

### Filamentation in mutant DHFR strains is not due to TLD and is reversible

A moderate level of thymine deficiency – thymine limitation – affects cell shape by increasing its diameter but not the length, making cells less rod-like, and certainly not filamentous (Pritchard & Zaritsky, 1970; Zaritsky & Pritchard, 1973). A more severe level of thymine deprivation causes, under certain conditions, thymineless death (TLD), a phenomenon accompanied by cell filamentation (Ahmad *et al*, 1998; Bazill, 1967; Sangurdekar *et al*., 2011; Zaritsky *et al*, 2006). Prior work (Sangurdekar *et al*., 2010; Sangurdekar *et al*., 2011) also showed that thymine limitation is metabolically different from thymineless death, as DNA damage and SOS induction happens only in the latter, and cell death happening due to erroneous DNA replication. The phenotypic, metabolic as well as expression signatures of mutant DHFR strains seemed therefore, to resemble a case of thymine deprivation and TLD. To find out if mutant strains represented a case of TLD, we assessed the viability of mutant strains at 30°C (permissive temperature) on solid media after several hours of growth at 42°C (restrictive temperature) in the presence of amino acids (filamentation condition). Sangurdekar *et al* (Sangurdekar *et al*., 2010) reported exponential loss in viability in TLD after one hour of growth under thymineless conditions. The idea was that if cells underwent TLD under conditions of filamentation, they would no longer be able to resume growth/form colonies at permissive temperature. To that end, we induced filamentous phenotype by incubating W133V mutant cells for 4 hours at 42°C, and then monitored cell recovery at room temperature (see *Methods*). Figure 4A shows a representative example of morphology of W133V filament that begins to undergo slow division initiated at the poles at permissive temperature. Most progeny cells of normal size appeared after 5-6 hours of growth at low temperature, indicating no loss in viability.

**Figure 4:**
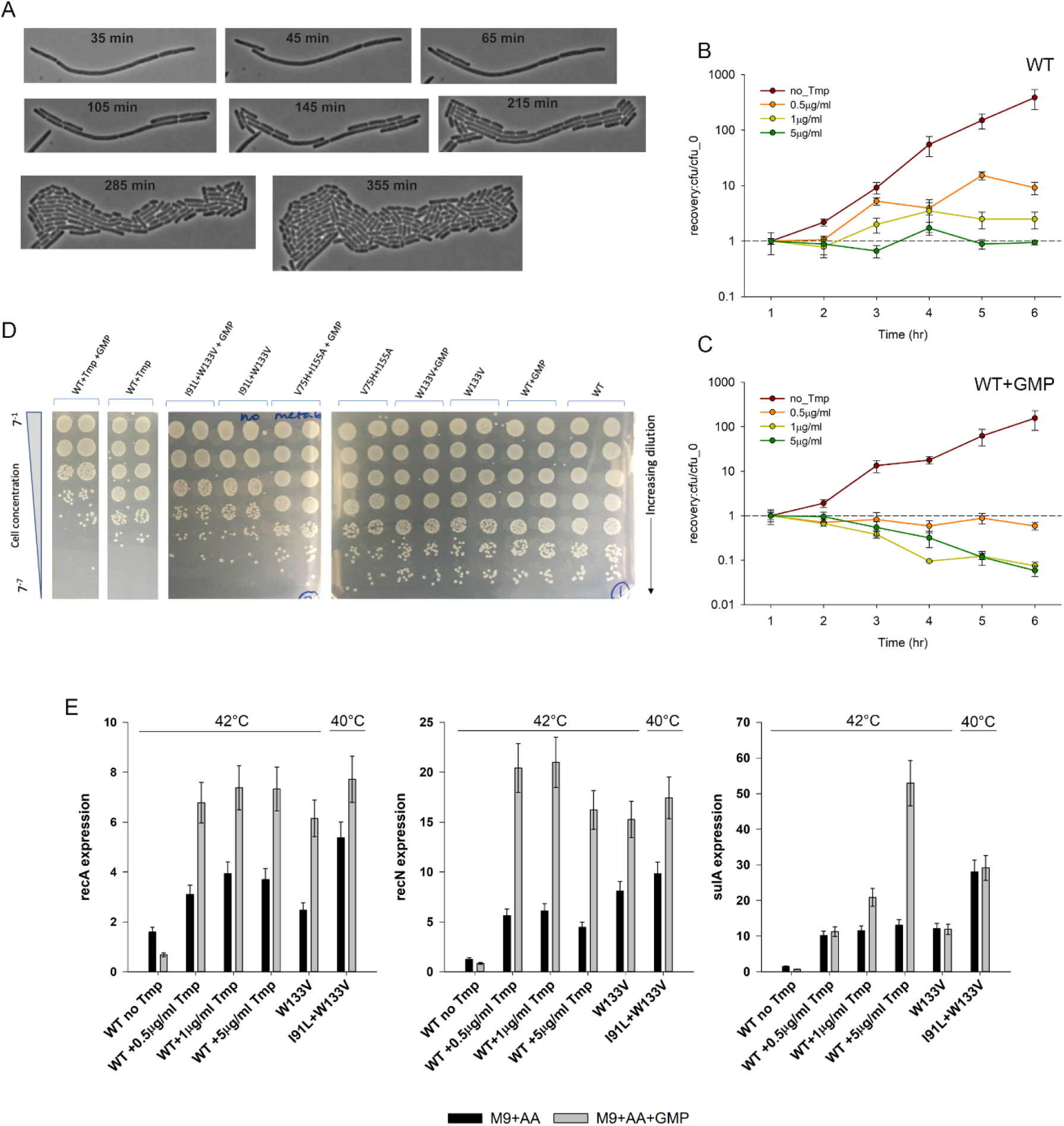
Filamentation in mutant DHFR strains is completely reversible and does not represent TLD. (A) Mutant W133V was grown in amino acid supplemented M9 medium (M9+AA) for 4 hours at 42°C, and subsequently placed on M9 agar pads and their growth was monitored at room temperature. Shown are phase contrast images taken from different time points throughout the time-lapse experiment. Unlike cells experiencing TLD, an irreversible phenomenon, W133V DHFR cells recover and resume growth at low temperature. (B and C) WT cells were treated with different concentrations of Tmp at 42°C for varying amounts of time in amino acid supplemented M9 medium (panel B) or in M9 media supplemented with both amino acids and GMP (panel C), following which they were spotted on M9+AA plates and allowed to grow at 30°C. Colonies were counted next day. In the presence of only amino acids, there was no loss in viability for any concentration of Tmp (panel B), despite extensive filamentation (Figure EV1E). In contrast, in the presence of amino acids and GMP, the cells showed sharp loss in viability when grown at high Tmp concentrations. In both panels, error bars represent SD of two biological replicates. The dashed line represents cfu at one-hour timepoint. (D) WT (with and without 5 μg/ml Tmp) and mutants were grown for 6 hours at 42°C in M9+AA medium without or with GMP, and subsequently diluted serially and spotted on M9+AA agar plates and allowed to grow at 30°C till visible colonies were formed. While WT treated with high Tmp shows loss in cfu, indicating TLD, no loss in viability was observed for WT or mutants. (E) Expression of SOS response genes recA, recN and sulA when mutant strains and Tmp treated WT cells are grown in M9 medium supplemented with amino acids (black bars) or in the presence of both amino acids and GMP (gray bars) at 42°C (I91L+ W133V was grown at 40°C). Error bars represent SD of 2-3 biological replicates.

Earlier studies have also shown that inhibition of DHFR activity by Trimethoprim (Tmp) under conditions of both amino acids and purine supplementation leads to TLD (Kwon *et al*., 2010). We found that when WT cells were grown on media supplemented with *both* amino acids and a purine source (GMP), they showed loss of colony forming units (indicating death) only when subjected to *very high* Tmp concentrations (Figure 4C), similar to (Kwon *et al*., 2010). On the other hand, supplementation with only amino acids was bacteriostatic under all conditions of Tmp (Figure 4B). Since mutant DHFR strains incur only *partial* loss of DHFR activity, they resemble lower concentrations of Tmp treatment, and therefore did not show any substantial loss in colony forming units even when grown in the presence of amino acids and GMP (Figure 4D), though the extent of SOS response increased substantially under this condition, indicating greater DNA damage (Figure 4E). Despite this, the fact that the mutant DHFR strains retained viability, reveals that TLD and cell survival depends crucially on the extent to which DHFR activity is compromised, a condition that is only achieved with lethal doses of Tmp. However, the data also shows expression of SOS genes alone cannot differentiate between the bacteriostatic and bactericidal regimes (Figure 4E). Collectively, these experiments disentangle the relationship between the regimes of DHFR activity and filamentation and TLD. This study also highlights the considerable overlap that exists between characteristics of thymine limitation and thymineless death and emphasizes the fact that these two conditions are not an all or none phenomenon.

### Supplementation of dTMP in the medium restores dTDP/dTTP levels and rescues filamentation

Since mutant strains had very low dTMP/dTTP levels, we grew WT and mutant strains in minimal medium that was supplemented with both amino acids and 1mM dTMP and carried out measurement of cell length as well as metabolomics analysis. Addition of dTMP largely rescued filamentation of mutant strains (Figure 5A). Metabolomics analyses showed that under this condition, I91L+W133V mutant had much higher levels of both dTDP and dTTP relative to WT (Figure 5B). Concentration of the other deoxyribonucleotides dATP/dCTP levels however, remained high despite addition of dTMP (Figure 5B). Therefore, we conclude that higher amounts of dTTP presumably reduce the imbalance in relative concentrations of deoxy-nucleotides, thus relieving DNA damage and filamentation (see above).

**Figure 5:**
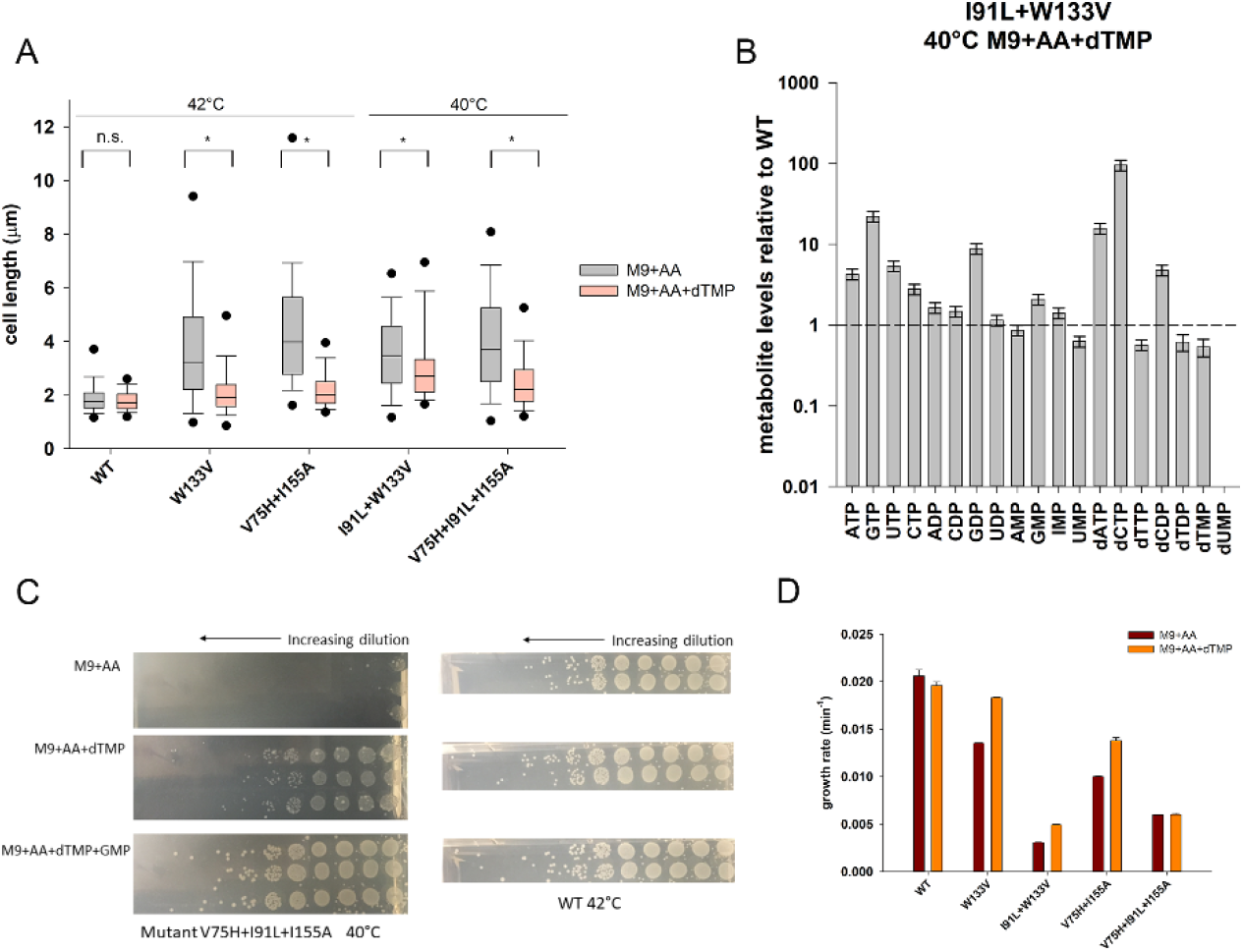
Supplementation of dTMP alleviates dTDP/dTTP levels and rescues filamentation. (A) Distribution of cell length of WT and mutant strains when grown in M9 medium supplemented with amino acids (gray) or with both amino acids and 1mM dTMP (pink). WT, W133V and V75H+I155A strains were grown at 42°C while I91L+W133V and V75H+I91L+I155A mutants were grown at 40°C. 1mM dTMP largely rescues filamentation of mutant strains (* indicates the median cell lengths were significantly different, Mann-Whitney test, p-value <0.001). The central band in the box plots represents the median of the distribution, the box ends represent the 25^th^ and 75^th^ percentile, the whiskers represent the 10^th^ and 90^th^ percentile, while the dots represent the 5^th^ and 95^th^ percentile. Data was usually obtained from 2-3 biological replicates. The number of cells used to derive the boxplot distributions in the different panels range usually between 150 to 300 (please refer to Figure 5 source data for exact number of cells for each dataset). (B) Abundances of selected nucleotides in I91L+W133V mutant when grown for 4 hours at 40°C in M9 medium supplemented with both amino acids and 1mM dTMP. Metabolite levels were normalized to those of WT grown under similar conditions. dTDP and dTTP levels recover and are now comparable to dTMP levels. Error bars represent SEM of 3 biological replicates. The dashed line in represents value of 1 for WT. (C) Mutant V75H+ I91L+I155A grows very poorly (in terms of colony forming units, cfu) on a minimal media agar plate supplemented with amino acids (M9+AA) at 40°C. Supplementation of additional dTMP increases the cfu by several orders at the same temperature, while supplementation with both pyrimidine (dTMP) and purine (GMP) allows it to grow as good as WT. In comparison, WT was grown at 42°C under different supplementation conditions. In all cases, cultures were 7-fold serially diluted for the next spot. The three rows (two rows for WT) in each condition represent biological replicates. (D) Comparison of growth rates of WT and mutant DHFR strains at 42°C (40°C for I91L+W133V and V75H+I91L+I155A) in minimal medium that is supplemented with amino acids and/or 1mM dTMP. Except for W133V and to a lesser extent for V75H+I155A, the effect of dTMP on growth rates is only modest. Error bars represent SEM of 3 biological replicates.

Moreover, supplementation of dTMP and amino acids also allowed mutants W133V, I91L+W133V and V75H+I91L+I155A to form higher counts of colony forming units (cfu) than with only amino acids at their respective filamentation temperatures (Figure 5C shows images for V75H+I91L+I155A at 40°C). However, we note that cfu count for mutants in the presence of dTMP were still orders of magnitude lower than those of WT, indicating that thymine only rescues cell length, and not growth defects. This was also supported by the absence of growth rate rescue with dTMP (Figure 5D).

### Low dTDP/dTMP ratio: possibility of inhibition of Thymidylate Kinase?

Interestingly, we found that while dTMP levels are low in mutant DHFR strains (10% and 50% of WT levels in I91L+W133V and W133V mutants respectively), dTDP levels were far lower (1% and 20%) (Figure 2C). On the other hand the ratios of dTDP to dTTP were approximately equivalent in both WT and mutants. Supplementation of dTMP in the medium however, restores the relative abundances of dTMP to dTDP to dTTP to approximately WT level in I91L+W133V (Figure 5B). It raises the possibility that the pyrimidine biosynthesis pathway enzyme Thymidylate Kinase (Tmk), which phosphorylates dTMP to dTDP, might be inhibited in mutant DHFR strains. Previous reports suggest that dUMP and dCTP can act as competitive inhibitors for Tmk (Nelson & Carter, 1969), and interestingly, we found that both dUMP and dCTP levels were highly upregulated in mutant cells (Figure 2C), due to inefficient conversion of dUMP to dTMP when DHFR activity was reduced (Figure EV2C). To find out if intracellular accumulation of dUMP and dCTP in mutant cells is sufficient to inhibit Tmk, we overexpressed and purified his-tagged Tmk from *E. coli* cells and tested the potential inhibitory effect of dUMP and dCTP *in vitro*. In comparison to its cognate substrate dTMP for which the K_M_ is 13μM (Figure 6A), the K_M_ for dUMP is 450μM (Figure EV3A), which indicates that dUMP is a much weaker substrate as compared to dTMP. The apparent K_I_ of dUMP for Tmk was 3.9mM (Figure EV3B). Given that dUMP concentration inside WT *E. coli* cells is 0.01mM (Møllgaard & Neuhard, 1983), its concentration inside mutant DHFR cells would be in the range of 0.5mM (50-fold higher levels), which is not enough to cause substantial inhibition of Tmk *in vivo*. We next carried out an activity assay of Tmk in the presence of varying amounts of dCTP. Depending on magnesium concentration, the K_I_ of dCTP ranged between 2-4 mM (Figure EV3C). Considering intracellular dCTP concentration in WT cells to be 35μM (Bennett *et al*., 2009), those in mutant cells are about 0.7mM (20-fold higher levels). Hence, like dUMP, intracellular dCTP levels are too low to show any substantial inhibition of Tmk activity *in vivo*. To conclude, though dUMP and dCTP have the potential to inhibit Tmk activity, their concentrations inside mutant cells are not high enough to achieve that.

**Figure 6:**
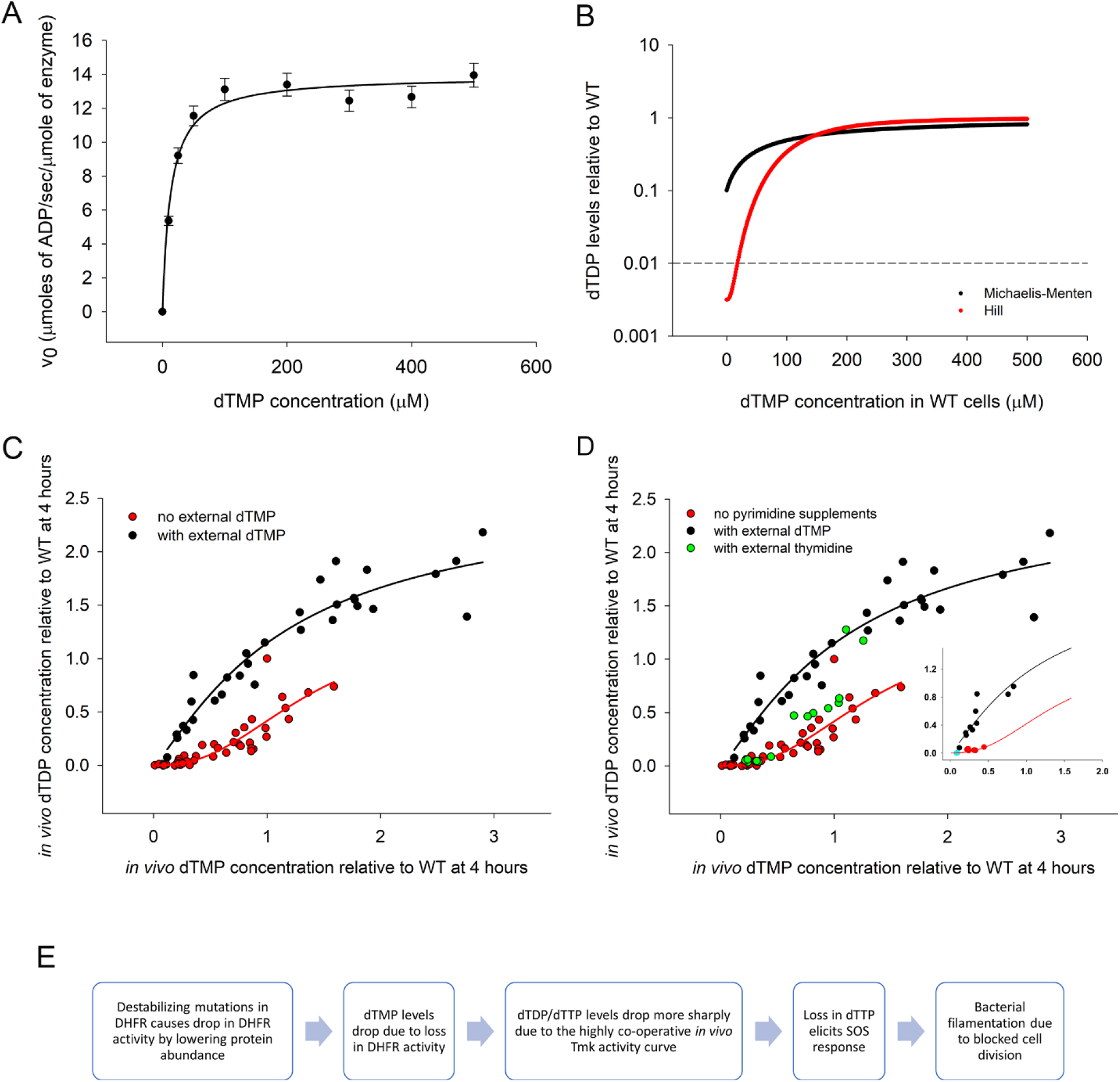
In vivo enzyme kinetics of Tmk is highly cooperative and different from that in vitro. (A) in vitro activity of purified Thymidylate Kinase (Tmk) as a function of dTMP concentration shows Michaelis-Menten (MM) like kinetics. ATP concentration is saturating at 1mM. The K_M_ of dTMP is 13μM. The error bars represent SD of three technical replicates. (B) For a 10-fold drop in intracellular dTMP concentration in mutant relative to WT (as seen in I91L+W133V mutant), we calculate dTDP levels in mutant (relative to WT) as a function of various assumed intracellular concentrations of dTMP in WT (shown along x-axis), as absolute value of this is not known experimentally, assuming MM kinetics (black line) as shown in panel (A) or Hill kinetics with coefficient of 2.5 (red line) as shown in panel (C). The dotted line corresponds to the experimentally observed dTDP ratio of 0.01 for I91L+W133V mutant. This ratio is realized only for the Hill-like curve. (C) Apparent in vivo activity kinetics of Tmk enzyme using steady state dTMP and dTDP levels obtained from metabolomics. The plot includes data from WT, mutants W133V and I91L+W133V, as well as WT treated with 0.5μg/ml Trimethoprim, obtained at different time points during growth. Data points represent metabolite levels for all individual biological replicates without averaging. The black data points were acquired during growth in the presence of different concentrations of external dTMP (0.25, 0.5, 1, 2 and 5mM), while red points were from conditions with no external dTMP. Both black and red solid lines represent fit to Hill equation. (D) The green points were acquired during growth of WT, W133V and I91L+W133V mutants in the presence of different concentrations of thymidine, which follow the red curve. The inset plot shows the same graph with selected datapoints for I91L+ W133V mutant. The cyan circle shows I91L+W133V mutant in the absence of any metabolite supplementation, while red and black points indicate metabolite levels following thymidine and dTMP supplementation respectively. (E) A flow chart summary of the actual chain of events triggered by destabilizing mutations in DHFR that eventually lead to bacterial filamentation, as revealed in this study.

### Cooperative enzymatic activity of Tmk in vivo explains low dTDP levels

As discussed previously, our metabolomics data shows that in mutant cells, dTDP levels fall far more precipitously than dTMP concentrations. If Tmk follows the same Michaelis-Menten (MM) like dependence on intracellular dTMP concentrations as seen *in vitro*, we find that dTDP levels would never drop as low as the experimentally observed value for mutants (Figure 6B, black line for I91L+W133V) for any assumed intracellular dTMP concentration for WT. To resolve this inconsistency, we attempted to elucidate the *in vivo* activity curve of Tmk enzyme from dTDP and dTMP levels measured from the metabolomics data for WT and mutant cells. To get a broad range of data, we measured metabolite levels for WT and several mutants at different time points during growth both in the absence and presence of different concentrations of external dTMP in the medium. Surprisingly, we found that the data points from our metabolomics experiments traced two different curves depending on whether there was external dTMP in the medium. Data points derived in the *absence* of external dTMP had a long lag, followed by a more cooperative increase (red points in Figure 6C), while the data points corresponding to added dTMP, appeared to follow the traditional MM curve (black data points in Figure 6C) and was similar to the enzyme activity of Tmk observed *in vitro*. We fitted both datasets to a Hill equation. For the conditions without added dTMP (red data points of Figure 6C) Hill-coefficient of 2.5 was obtained suggesting strong positive cooperativity. On the other hand, when dTMP was added, the dTDP vs dTMP curve (black datapoints in Figure 6C) was fitted with a Hill coefficient of 1.2. However, the fit was not significantly different from the traditional MM model (p-value = 0.36). Based on the Hill-like curve in Figure 6C, we can say that for intracellular dTMP concentrations below 10μM, a 10% drop in dTMP levels (as seen for I91L+W133V) would cause dTDP levels to drop to 1% or less (Figure 6B, red line), largely because of the long lag. Therefore, a Hill-like dependence of Tmk activity on dTMP concentrations can explain the disproportionately low dTDP levels in mutant strains.

However, we note that metabolomics does not directly report on the kinetics of an enzyme *in vivo*. Rather, it reports on the steady state levels of metabolites present inside the cell at any given time. Moreover, unlike *in vitro* activity measurement conditions where an enzyme functions in isolation, *in vivo* Tmk is a part of the pyrimidine biosynthesis pathway that involves several sequentially acting enzymes as follows:

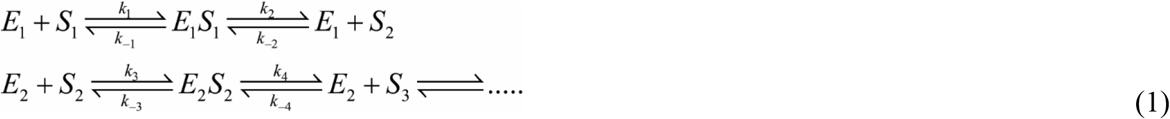

Where all metabolites (*S*_1_, *S*_2_, *S*_3_,..) are at steady state.

To shed light on whether steady state metabolomics data can give us insight into the kinetics of enzymes *in vivo*, we carry out a simple exercise as follows: for a system (like equation (1)) where a series of enzymes work sequentially, we assume two scenarios.

<u>Case 1</u>, we assume that each enzyme in this pathway follows Michaelis-Menten kinetics and that the *product* of each enzyme has very low affinity to bind the enzyme back. Using the detailed derivation in the Supplementary Text, we arrive at the following equation:

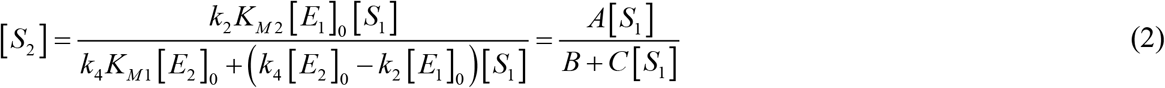

Where 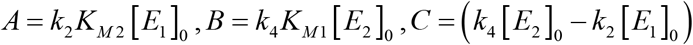 and *K*_*M* 1_ and *K*_*M* 2_ are the Michaelis constants of enzymes *E*1 and *E*2 for *S*1 and *S*2, [*E_i_*]_0_ is a total concentrations (i.e. free+bound) of an i-th enzyme. Equation (2) tells us that if individual enzymes follow MM kinetics (assumptions of Case 1), then we expect that the steady state concentration of any product in the pathway will also follow a hyperbolic or MM like dependence on its substrate concentration.

<u>Case 2</u>, we assume that *E*1 or *E*2 or both follow Hill-like kinetics (due to reasons we elaborate in the Discussion) in the following form:

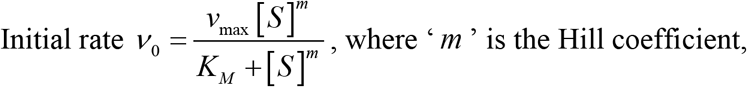

then for the pathway of enzymes at steady state (as in Eq (1)), *S*2 has the following dependence on *S*1:

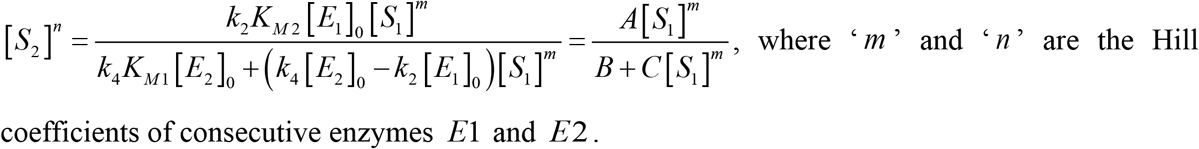

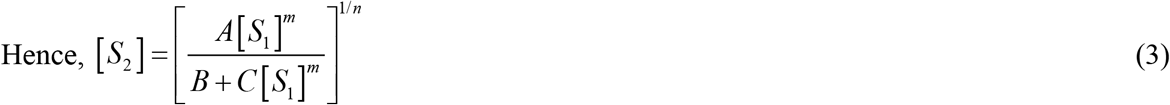

As shown in Supplementary Text, Equation (3) tells us in Case 2 scenario, *S*2 will show a Hill like dependence on *S*1 with positive cooperativity with the assumption that *m* > *n*.

The above analyses with both Case 1 and 2 suggest that the dependence of the steady state concentrations of metabolites on each other along a metabolic pathway is reflective of the kinetics of the concerned enzyme *in vivo*. In other words, since the Tmk activity curve *in vivo* is Hill-like in the *absence* of external dTMP, and MM like in the presence of added dTMP (Figure 6C), it is reasonable to infer that *kinetics* of Tmk *in vivo* is Hill-like without added dTMP and MM like in the presence of external dTMP. Of course, this analysis assumes that ATP, the second substrate of Tmk is not limiting, and therefore kinetics of Tmk is pseudo-unimolecular with respect to dTMP. Indeed, our metabolomics data validates the assumption of pseudo-unimolecular in vivo kinetics of Tmk by showing that even in mutant DHFR strains, ATP is not depleted, hence not limiting.

### Supplementation of thymidine retains cooperative behavior of Tmk in vivo

dTMP, the substrate of Tmk, comes from two different sources inside the cell: the *de novo* pyrimidine biosynthesis pathway through conversion of dUMP to dTMP by thymidylate synthase, and the pyrimidine salvage pathway through conversion of thymine to thymidine to dTMP. Mutant DHFR strains which are unable to efficiently convert dUMP to dTMP through the *de novo* pathway due to reduced folate activity, rely substantially on the salvage pathway for their dTMP supply, as has been shown for catalytically inactive mutants of DHFR (Rodrigues & Shakhnovich, 2019). Hence, we next asked the question: what happens to the Tmk activity curve *in vivo* if dTMP is produced (largely) through the salvage pathway instead of being directly supplied from an external source? To that end, we supplemented the growth medium with intermediates from the salvage pathway, namely thymine and thymidine. While supplementation of thymidine increased intracellular dTMP levels for WT as well as mutants (Figure EV4A), the dTDP vs dTMP levels followed the Hill like curve (Figure 6D, inset figure shows data for the I91L+W133V mutant in the absence of supplementation and in the presence of added thymidine and dTMP). This shows that direct supplementation of the substrate (dTMP) of Tmk results in very different enzyme kinetics compared to when a precursor of dTMP is supplied externally.

Quite surprisingly and contrary to WT, *mutant* strains did not use external thymine base towards increasing intracellular dTMP (Figure EV4A), though it was up taken by the cells (Figure EV4B). We ruled out inhibition of DeoA enzyme (which interconverts thymine and thymidine) in mutants, as thymidine supplementation increases thymine levels significantly (Figure EV4C). However, I91L+W133V mutant had considerably lower level of deoxy-D-ribose-1-phosphate (dR-1P) (Figure EV4D) which is used as a substrate by the enzyme DeoA to synthesize thymidine from thymine. This strain also accumulated large excess of deoxy-D-ribose-5-phosphate (dR-5P) (Figure EV4D), indicating that isomerization of the sugar dR-1P to dR-5P through DeoB enzyme might be one of the reasons for the lack of thymine utilization. This scenario is supported by the recent finding that cells that evolved to grow on small amounts of thymine supplement on the background of inactive DHFR mutationally deactivated DeoB thus blocking the channeling of dR-5P towards glycolysis and providing sufficient amount of dR-1P towards thymidine synthesis in the salvage pathway (Rodrigues & Shakhnovich, 2019).

### Is cooperativity a general signature of enzyme activity in vivo?

Since *in vivo* kinetics of Thymidylate Kinase is cooperative and Hill-like, we next sought to find out if this is a general characteristic of metabolic enzymes *in vivo*. To that end, we looked at the enzyme Adenylate Kinase (AK), which is an essential enzyme in *E. coli* that interconverts adenylate currencies AMP, ADP and ATP. Importantly, from a previous study (Adkar *et al*., 2019), we had metabolomics data available for several stabilizing and destabilizing mutants of AK in the absence and presence of external AMP. This provided us a handle to span a large range of intracellular AMP and ADP concentrations, in turn allowing us to get an insight into the kinetics of AK enzyme *in vivo*. When we overlay the relative AMP and ADP concentrations for all mutants (Figure 7A), we find that much unlike Tmk, the *in vivo* activity curves of AK obey Michaelis-Menten like kinetics both in the presence and absence of external AMP supplementation.

**Figure 7:**
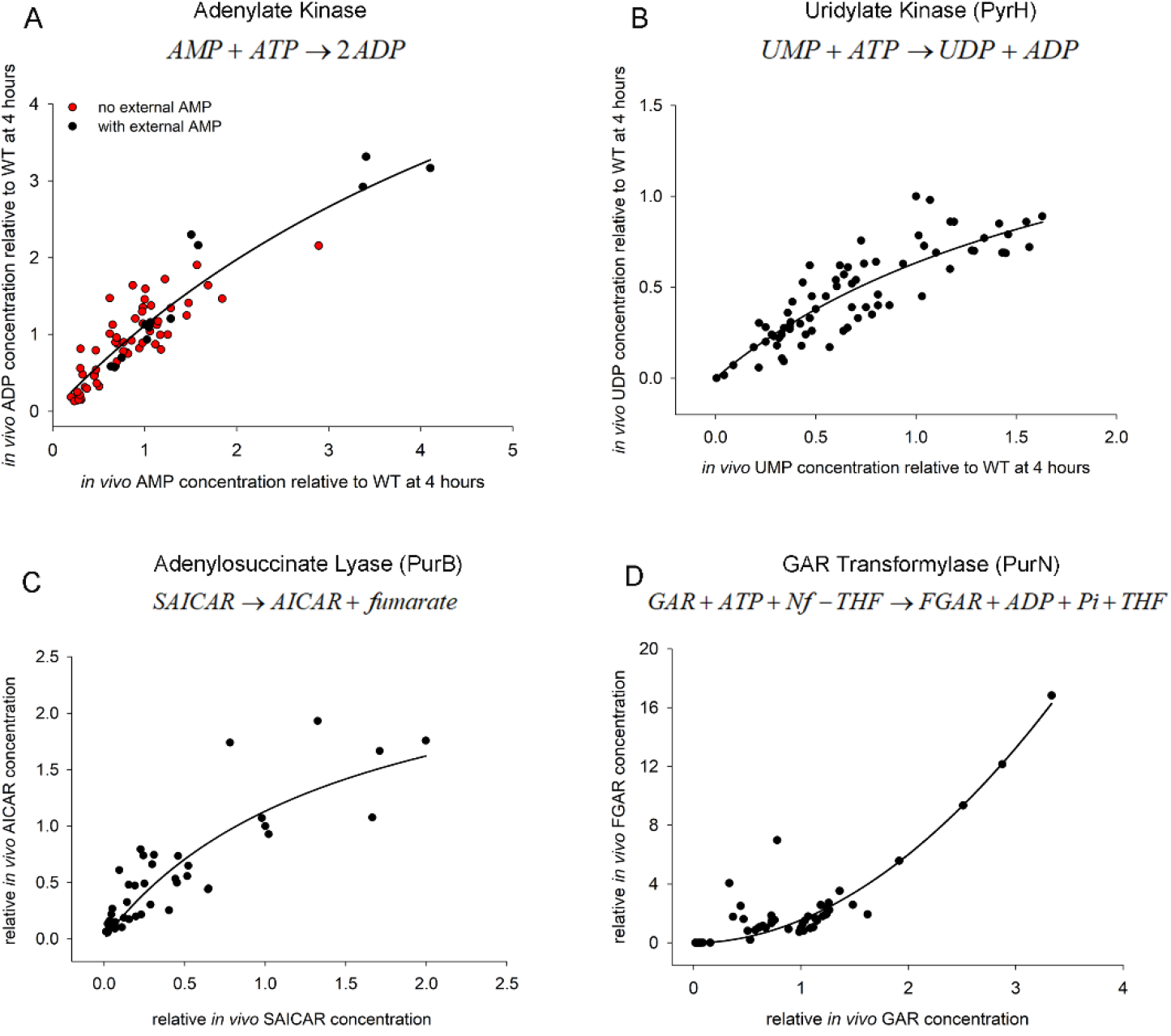
In vivo activity curves of representative E. coli enzymes show hyperbolic or cooperative behavior. Steady state levels of different metabolites obtained from metabolomics were used to derive activity curves for (A) Adenylate Kinase (AK) (B) Uridylate Kinase (PyrH) (C) Adenylosuccinate Lyase (PurB) and (D) GAR Transformylase (PurN). AK, PyrH and PurB appear to follow hyperbolic MM kinetics in vivo, while PurN shows Hill-like dependence with cooperativity. In panel (A), the red data points were acquired during growth in amino acid supplemented M9 medium in the absence of external AMP and are derived from WT and various stabilizing and destabilizing point mutations of AK (Adkar et al., 2019) as well as DHFR mutants used in the present study, while black points were derived exclusively from cultures of AK mutants supplemented with 1mM external AMP (Adkar et al., 2019). Data points for all other panels (B-D) are derived from mutant DHFR strains in the present study. Since SAICAR, AICAR, GAR and FGAR levels were undetectable in WT, all data points in (C-D) are relative to those observed in I91L+W133V mutant following 4 hours of growth at 40°C. Data points in all panels represent metabolite levels for individual biological replicates without averaging.

Intrigued by the AK data, we set out to look for similar data for other *E. coli* enzymes. However, in the absence of direct mutational data for any other enzyme, it is difficult to obtain a large enough dynamic range of intracellular metabolite concentrations. In case of Tmk, it was possible to obtain large variations in dTMP levels as drop in folate end products due to mutations in DHFR directly affect dTMP levels. However, we reasoned that since folate products are utilized in both *de novo* purine and pyrimidine biosynthesis pathways, several intermediates in those pathways might show varying levels of accumulation. Due to inherent low stability of certain intermediates, we could only detect a handful of metabolites with confidence. However even that allowed us, using our metabolomics data, to construct activity curves for three more enzymes: Uridylate Kinase (PyrH) from the *de novo* pyrimidine biosynthesis pathway, and Adenylosuccinate Lyase (PurB) and GAR Transformylase (PurN) from the *de novo* purine biosynthesis pathway. As shown in Figure 7B, C, PyrH and PurB show a hyperbolic, Michaelis-Menten-like dependence of product on substrate levels, like Adenylate Kinase. Strikingly, PurN (Figure 7D) shows a long lag followed by steep transition and fitted Hill-like kinetics with high significance (p-value <0.0001 for MM as null model), much like Tmk, with an estimated Hill coefficient of ~2. However, since the substrate GAR (Glycineamide Ribonucleotide) for this enzyme is not commercially available, we were unable to test the kinetics of PurN in the presence of GAR supplementation.

We therefore conclude that while enzymes like Tmk and PurN that show Hill-like kinetics *in vivo*, there are many that retain traditional MM kinetics in the cell. Presumably, this effect is dictated by the complex cellular milieu, on which we elaborate more in the Discussion.

## Discussion

Metabolic networks of cells are inherently intertwined, with substrates and products of one pathway being utilized by another pathway. As a result, perturbations produced in one pathway can easily percolate into others, usually magnifying the effects. The folate pathway or the 1-carbon metabolism pathway is a classic example of this, as reduced folates act as 1-carbon donors during biosynthesis of purines, pyrimidines and amino acids. Kwon et al (Kwon *et al*, 2008) showed that for inhibition of DHFR activity using trimethoprim, accumulation of substrate dihydrofolate (DHF), in turn, results in inhibition of another downstream enzyme critical to folate metabolism: folylpoly-gamma-glutamate synthetase (FP-gamma-GS), in a domino like effect (falling DHFR activity triggers a fall in the other enzyme’s activity too). In this work, we show that in *E. coli* strains that harbor destabilizing mutations in *folA* gene, reduced DHFR activity strongly affects, among other factors, the pyrimidine biosynthesis pathway by reducing production of dTMP from dUMP via thymidylate synthase (ThyA) that uses a derivative of THF as one carbon source. Much like a domino effect, such drop in dTMP levels due to mutations in DHFR results in a precipitous drop in dTDP/dTTP, mainly due to the strong cooperative *in vivo* activity of another downstream essential enzyme Thymidylate Kinase (Tmk) in the pyrimidine biosynthesis pathway. Drop in dTTP level eventually leads to an imbalance in the levels of deoxynucleotides, presumably causing errors in DNA replication, SOS response and filamentation due to blocked cell division (Figure 6E).

An important finding from the current study is that enzymes can exhibit different kinetics *in vivo* depending on the source of the substrate. In case of Tmk, the enzyme showed a conventional Michaelis-Menten type *in vivo* activity when dTMP was externally supplied through the growth medium. However, when an equivalent concentration of dTMP was produced by the cell itself using its own cascade of enzymes in the pathway, it showed a dramatically different cooperative (Hill-like) activity. There are two important questions that arise out of this observation: first, why is the intrinsic *in vivo* activity of Tmk Hill-like? Second, what causes this shift from Hill-like to Michaelis-Menten (MM)? One of the most straightforward reasons for Hill-like enzyme activity is allosteric substrate binding. However, purified Tmk *in vitro* shows perfect MM kinetics, ruling out any intrinsic allostery of the enzyme. We also found that even in the presence of high concentrations of dUMP and dCTP (the known inhibitors of Tmk), the activity of Tmk conforms to MM kinetics ruling out these metabolites as allosteric regulators (Figure EV3D). There are several other metabolites (GTP, GDP, GMP, dATP, UTP) in our targeted metabolomics analyses that are highly upregulated in the mutant strains (Figure 2C), however they were upregulated in the presence of external dTMP as well (condition with MM like kinetics) (Figure 5B), thereby ruling out their role in allosteric modulation of Tmk activity. The other possible mechanism of Hill-like kinetics is ‘limited diffusion’ of one or more of the interacting components of a reaction (Savageau, 1995, 1998). Conventional MM enzyme kinetics relies on the assumption of free diffusion, and hence laws of mass action are obeyed. However, in case of limited diffusion, conditions of spatial uniformity are no longer maintained; hence, law of mass action is not applicable. Theoretical work as well as simulations (Frank, 2013; Kopelman, 1988; Li *et al*, 1990; Liebovitch *et al*, 1987; Savageau, 1995, 1998) have shown that such diffusion limited reactions often exhibit kinetics with Hill-like coefficients that are significantly higher than 1, and often fractional (so called fractal kinetics), similar to Hill coefficients obtained with our data (Figure 7C, Hill coefficient=2.5). It has also been postulated that biological reactions, especially those that happen in dimensionally restricted environments like 1D channels or 2D membranes exhibit fractal like kinetics (Liebovitch *et al*., 1987; Schnell & Turner, 2004). In our case, it seems more reasonable that it is the substrate dTMP that has limited diffusion rather than the enzyme Tmk itself, since addition of external dTMP alleviates the Hill-like effect (in Supplementary Text, we show a derivation of Hill-like enzyme kinetics assuming that only the substrate is diffusion limited, using a power law formalism as developed by Savageau (Savageau, 1995)). But why should dTMP be diffusion limited? Substantial work in the recent past has shown that metabolic enzymes of a pathway, including those involved in purine biosynthesis (An *et al*, 2008; French *et al*, 2016) as well as in 1-carbon metabolism (Bhattacharyya *et al*., 2016) form a metabolon, a supramolecular complex comprised of transiently interacting enzymes, that allows efficient channeling of metabolites. Though channels help in easy exchange of metabolites between consecutive enzymes and prevent their unwanted degradation or toxicity in the cytosol, they have reduced dimensionality compared to the cytosol, thereby making motion less ‘random’ and hence limiting diffusion of the substrate/products. In our case, it is possible that enzymes of the pyrimidine biosynthesis pathway as well as the salvage pathway form a metabolon, that limits diffusion of dTMP. External dTMP on the other hand, is free to diffuse in the cytoplasm, and hence results in traditional MM kinetics to emerge with a Hill coefficient of 1.

Though we do not have direct evidence of Tmk being involved in a metabolon, our study does show some circumstantial evidence. Our previous work on DHFR showed that toxicity and filamentation upon over-expression might be a hallmark of metabolon proteins (Bhattacharyya *et al*., 2016), through sequestration of neighboring/sequential proteins in the pathway. On a similar note, we found that overexpression of Tmk in WT *E. coli* cells is toxic and lead to filamentation (Figure EV5A, B), strongly suggesting that Tmk might be part of a metabolon. It is worth mentioning at this point that Tmk overexpression in mutant DHFR cells does not rescue filamentation (Figure EV5A). This is consistent with the metabolon hypothesis, since dTMP produced by the cell would still be confined to the metabolon, and hence overexpressed Tmk cannot overcome the problem of diffusion limitation of dTMP.

PurN is another enzyme that shows strong Hill-like kinetics in our study, and it is worth noting that the mammalian analog of PurN (the GAR Transformylase) constitutes one of the core proteins of the purinosome (Deng *et al*, 2012). Though in the absence of direct metabolic complementation studies, it is difficult to pinpoint which metabolite might be diffusion limited, our study nevertheless strongly points towards the existence of a similar purinosome complex in *E. coli* as well. This is also indirectly supported by the fact that overexpression of PurN is highly toxic in *E. coli* (Kitagawa *et al*, 2005). Regarding Adenylate Kinase (AK), though several enzymes use up ATP, AMP and ADP, AK as such is not part of a pathway, and which might be the reason behind lack of any substantial cooperativity *in vivo*. Previous data from co-IP experiments also showed no interaction with other *E. coli* proteins (Bhattacharyya *et al*., 2016). Moreover, data from the ASKA library as well as our own (Figure EV5, (Bhattacharyya *et al*., 2016; Kitagawa *et al*., 2005)) show that over-expression of AK is not toxic, again supporting non-involvement of AK in any kind of metabolon. Of course, not every enzyme in a pathway is part of a metabolon, as presumably exemplified by data from PyrH and PurB which are respectively parts of the *de novo* pyrimidine and purine biosynthesis pathways yet show hyperbolic *in vivo* kinetics. Though PurB is part of the mammalian purinosome, it is not part of the core complex, and is known to associate with weak interaction (Deng *et al*., 2012). Data from ASKA library show that PyrH is neutral upon over-expression (Kitagawa *et al*., 2005), though PurB is mildly toxic. PurB however catalyzes a second reaction outside the purine biosynthesis pathway, which might contribute to the observed toxicity.

For decades, enzyme activity has been studied *in vitro* with purified enzyme in dilute solution with excess substrate. Though *in vitro* measured parameters have been largely successful to interpret cellular data (Adkar *et al*., 2017; Rodrigues *et al*., 2016), in other cases they have only provided limited information (van Eunen *et al*, 2012). Substantial efforts in the recent past have therefore been directed towards replicating *in vivo* like conditions with purified enzymes (Davidi *et al*, 2016; Garcia-Contreras *et al*, 2012; Zotter *et al*, 2017). These include macromolecular crowding, pH conditions and buffer capacity (van Eunen, 2014). In this work, we present a potentially powerful general method for detecting *in vivo* activity of an enzyme using product to substrate ratios obtained from metabolomics and show how the *in vivo* activity curves of an enzyme can be markedly different (in case of Tmk strongly Hill-like) from its perfect MM like kinetics *in vitro*. Based on available literature and some of our preliminary experiments/observations on Tmk and other proteins, metabolon formation and subsequent diffusion limitation of one of the substrates (dTMP in case of Tmk) seems like the most probable mechanism. Future work will prove or disprove this hypothesis. However, regardless of the mechanism, this work provides convincing evidence that the cellular environment can modulate enzyme activity in a very fundamental way, which explains a key bacterial phenotype in our case.

In this study, we used metabolomics as the key tool to link molecular effects of mutations to phenotype and illustrate precise biochemical and biophysical mechanisms through which altered metabolite levels modulate bacterial phenotypes, in this case, filamentation. Detailed metabolomics analysis allowed us to pinpoint the pathway and specific enzyme responsible for the phenotype and, surprisingly, it turned out to be far downstream from the mutant locus (folA). Furthermore, the culprit, Tmk, does not use products of the folate pathway as a cofactor. Nevertheless, it appears that perturbation of the folate pathway caused by mutations in DHFR propagated downstream in a domino-like manner to create a bottleneck in a specific metabolite dTDP triggering cellular SOS response and pronounced phenotypic effects manifested in altered cell morphology. Altogether our results show how metabolomics can be used as a stepping-stone from biophysical analysis of variation of molecular properties of enzymes to phenotypic manifestation of mutations and close the gap in the multi-scale genotype-phenotype relationship.

## Acknowledgements

This work was supported by NIH grant GM068670 to E.I.S.

## Author contributions

Conceptualization, S.Bh., S.Be., and E.I.S.; Methodology, S.Bh., S.Be., B.V.A., J.W., and E.I.S.; Formal Analysis, S.Bh., S.Be., B.V.A., and E.I.S.; Investigation, S.Bh., S.Be, B.V.A., J.W.; Writing-original draft, S.Bh., and E.I.S.; writing-review & editing, S.Bh., S.Be., B.V.A., and E.I.S.; Supervision, E.I.S.; Funding acquisition, E.I.S.

## Methods

### Strains and media

The mutant DHFR strains chosen for this study (W133V, I91L+W133V, V75H+I155A and V75H+I91L+I155A) were a subset of strains generated in and described in Bershtein *et al* (Bershtein *et al*., 2012). Briefly, using structural and sequence analyses, positions were chosen that were buried in the protein and located at least 4Å away from the active site, so that mutations introduced at these positions would have minimal effect on enzymatic activity. The identity of the mutations was selected such that they are appear with very low frequency in a multiple sequence alignment, and hence were intended to destabilize the protein, as was confirmed by stability measurements of the purified proteins (see Table below). The single mutants were in most cases mild to moderately destabilizing, and hence certain mutations were combined to increase the range of destabilization achieved. The following table summarizes the results of stability measurements on the single mutants (from (Bershtein *et al*., 2012)). However, the multiple site mutants could not be purified to quantities that were sufficient for *in vitro* biophysical and biochemical characterization.

| Mutant | Interaction | T<sub>m</sub> (°C) | ΔG (kcal/mol) | ΔΔG (kcal/mol) |
| --- | --- | --- | --- | --- |
| WT |  | 51.7 | -4.4 | 0 |
| V75H | Steric clash | 40.6 | -2.18 | 2.22 |
| I91L | Steric clash | 41.4 | -2.69 | 1.71 |
| W133V | Cavity formation due to large to small substitution | ND | -1.53 | 2.87 |
| I155A | Cavity formation due to large to small substitution | 38.5 | -2.42 | 1.98 |

Single as well as multiple site mutations were eventually introduced into the chromosomal copy of the *folA* at its endogenous locus keeping its regulatory region intact, and the effect of the mutations on its growth and morphology were measured at 30, 37 and 42°C. While WT, W133V and V75H+I155A could grow up to 42°C, mutants I91L+W133V and V75H+I91L+I155A could only survive up to 40°C, and hence these strains were grown at 40°C instead of 42°C. The reason for choosing a wide range of temperature instead of following a single conventional temperature of 37°C was that *E. coli* is a gut bacterium and inhabit hosts whose core body temperatures span a large range (37-38°C for mammals, 40-45°C for birds (Guenther *et al*, 2010; Torre-Bueno, 1976)). Moreover, since the chosen mutants were temperature sensitive, the phenotypic manifestation of the mutations was the largest at the extremes of temperatures, in case of *E. coli* at 42°C.

Wherever mentioned, M9 minimal medium *without amino acids* was only supplemented with 0.2% glucose and 1mM MgSO_4_ while M9 media *with amino acids* was supplemented with 0.2% glucose, 1mM MgSO_4_, 0.1% casamino acids, and 0.5 μg/ml thiamine. Casamino acids is a commercially available mixture of all amino acids except tryptophan, and cysteine is present in a very small amount. Wherever mentioned, 1mM GMP was used as a source of purine, while dTMP (thymidine monophosphate) was used at a concentration of 0.25-1mM. Thymidine was used at two different concentrations of 0.5mM and 1mM.

### Growth conditions

All strains were grown overnight from a single colony at 30°C, and subsequently the culture was diluted to a final OD_600_ of 0.01 in the specified medium and allowed to grow for 16-18 hours in Bioscreen C (Growth Curves, USA) at 30°C, 37°C or 42°C. Growth curves were fit to a 4-parameter Gompertz equation as described in (Bhattacharyya *et al*, 2017) to derive growth parameters. Error bars were calculated as SEM of three biological replicates.

### Light microscopy

Cells were grown overnight at 30°C from a single colony in the specified medium, diluted 1/100, and grown at various temperatures for 4 hours. For DIC images in Figures 1C, 1D, Figure EV1A, cells were pelleted, washed with PBS, and concentrated. DAPI staining (Molecular probes) was performed for 10 min at RT according to manufacturer instructions. 1 μl of a concentrated culture was then mounted on a slide and slightly pressed by a cover slip. DIC and DAPI images were obtained at room temperature by Nikon Ti Eclipse Microscope equipped with iXon EMCCD camera (Andor Technologies). For live phase contrast images and time-lapse experiments (Figure 4A), cells were mounted on supplemented M9 + 1.5% low melting agarose (Calbiochem) pads. Pads were then flipped on #1.5 glass dish (Willco Wells), and the images were acquired at room temperature with Zeiss Cell Observer microscope. For DIC images in Figure EV5A, cells were placed on agar pads and images were acquired with Zeiss Cell Discoverer microscope.

### Analysis of cell lengths

MicrobeTracker Suite [http://microbetracker.org/](Sliusarenko *et al*, 2011) was used to obtain distributions of cell length for phase contrast images and Zeiss Intellesis Module was used to analyze DIC images. On average, 500 cells were analyzed for each presented distribution. The cell lengths are represented in all figures as box-plots, where the boundaries of the box represent the 25^th^ and 75^th^ percentile, the line inside the box represents the median of the distribution, whiskers represent the 10^th^ and 90^th^ percentile, while the dots represent the 5^th^ and 95^th^ percentile.

### Statistical analysis

In our experiments, cell lengths of *E. coli* were not normally distributed (Shapiro-Wilk test). Hence non-parametric Mann-Whitney test was used to determine if the median cell lengths of two samples were significantly different. In Figure 6C, fits to two different models, Michaelis-Menten and 3-parameter Hill were compared using extra sum-of-squares F-test using GraphPad Prism software v9.0.0.

### Metabolomics

Cells were grown overnight at 30°C from a single colony in the specified medium, diluted 1/100, and re-grown. WT, WT+0.5μg/ml Tmp and W133V mutant were grown at 42°C, while mutant I91L+W133V was grown at 40°C. For time course experiment, aliquots were removed after 2, 4, 6 and 8 hours, and metabolites were extracted as described in (Bhattacharyya *et al*., 2016). Briefly, the cells were washed 2 times with chilled 1×M9 salts, and metabolites were extracted using 300μl of 80:20 ratio of methanol:water that had been pre-chilled on dry ice. The cell suspension was immediately frozen in liquid nitrogen followed by a brief thawing (for 30 seconds) in a water bath maintained at 25°C and centrifugation at 4°C at maximum speed for 10 minutes. The supernatant was collected and stored on dry ice. This process of extraction of metabolite was repeated two more times. The final 900μl extract was spun down one more time and the supernatant was stored in −80°C till used for mass spectrometry. Metabolite levels were averaged over 2-3 biological replicates as increasing sample size beyond 3 did not significantly change the SEM. In Figures 6 and 7, data points represent metabolite levels for all biological replicates without averaging.

### Expression of SOS response genes by qPCR

Cells were grown overnight at 30°C from a single colony in the specified medium, diluted 1/100, and grown at 37°C or 42°C for 4 hours. Based on OD_600_ of the cultures, a volume equivalent to 5×10^8^ cells were spun down (assuming 1 OD_600_=8×10^8^ cells) and Protect Bacteria RNA Mini Kit (Qiagen) was used to extract total RNA as described in (Bhattacharyya *et al*., 2018). Following reverse transcription (Bhattacharyya *et al*., 2018), expression of *recA, recN* and *sulA* genes were quantified using QuantiTect SYBR Green PCR kit (Qiagen) using the following primers:

recA_fwd ACAAACAGAAAGCGTTGGCG
recA_rev AGCGCGATATCCAGTGAAAG
recN_fwd TTGGCACAACTGACCATCAG
recN_rev GACCACCGAGACAAAGAC
sulA_fwd GTACACTTCAGGCTATGCAC
sulA_rev GCAACAGTAGAAGTTGCGTC

As it was difficult to find a reference gene that would be expressed to similar levels in WT vs mutant DHFR strains, we used total RNA to normalize the expression levels.

Expression levels reported are average of 3 biological replicates. Error bars in Figure 4 represent 12% of the mean value.

### Tmk protein purification

The tmk gene was cloned in pET28a plasmid between *NdeI* and *XhoI* sites with an N-terminal histag. BL21(DE3) cells transformed with the plasmid were grown in Luria Broth at 37°C till an OD of 0.6, induced using 1mM IPTG and grown for an additional 5 hours at 37°C. The protein was purified using Ni-NTA affinity columns (Qiagen) and subsequently purified by gel filtration using a HiLoad Superdex 75 pg column (GE). The protein was concentrated and stored in 10 mM potassium phosphate buffer (pH 7.2). The concentration of the proteins was measured by BCA assay (ThermoScientific) with BSA as standard.

### Tmk activity assay

Tmk catalyzes the following reaction 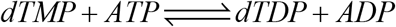, and the activity assay was carried out using the spectrophotometric assay as described in (Nelson & Carter, 1969). Briefly, the reaction mixture contained 5mM MgCl_2_, 65mM KCl, 350uM phosphoenolpyruvate (PEP), and 300uM NADH. To obtain K_M_ for dTMP, ATP concentration was fixed at 1mM, while dTMP concentration was varied from 10μM to 500μM. The reaction mix without enzymes was incubated at 25°C for 5 minutes, and the reaction was initiated by adding 100nM Tmk (final concentration) and 2 units of pyruvate/lactate dehydrogenase. The kinetic traces were recorded for every 5 seconds for a total time of 1 minute. The data corresponding to the first 20 seconds were fitted to a linear model to obtain initial rates. To obtain K_I_ of dCTP for Tmk, ATP and dTMP concentrations were fixed at 100μM and 1mM respectively, while dCTP concentration was varied from 0.5mM to 7.5mM. Since conversion of dUMP to dTDP also produces ADP, the K_I_ of dUMP could not be estimated by the spectrophotometric method. Instead, dTDP amounts produced in the reaction were determined by LC-MS. For the reaction, ATP and dTMP concentrations were fixed at 1mM and 100μM respectively, while dUMP concentration was varied from 0.25mM to 5mM. The reaction was quenched at 40 seconds using 80% MeOH. The resulting samples were subjected to LC-MS analysis to obtain dTDP levels. For Figure EV3(B and D), LC followed by mass spectrometry was used to directly measure dTDP levels.

#### Derivation of the relationship between steady state metabolite concentrations and kinetics of the enzyme *in vivo*

##### Case I: Sequential enzymes in pathway with Michaelis-Menten kinetics

The following is an example of a pathway where several enzymes work sequentially:

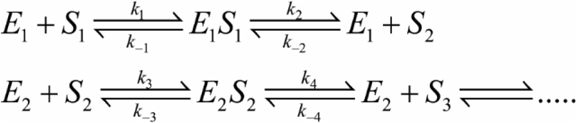

For the pathway at steady state, the concentrations of the reactants, products and intermediates do not change with time. Therefore,

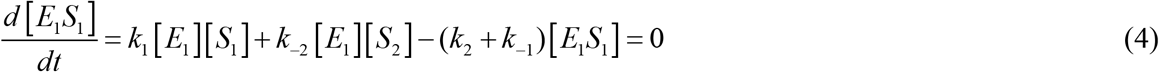

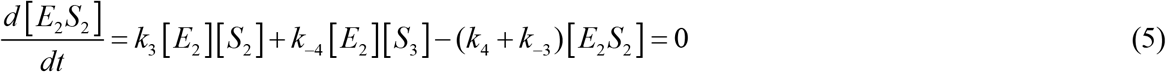

The enzyme concentrations can be written as:

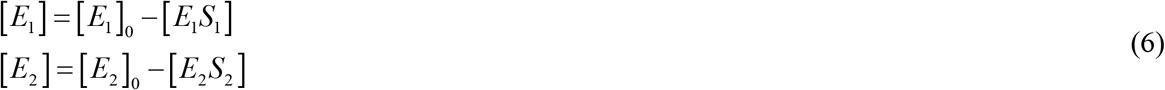

[*S*_1_], [*S*_2_] are the steady state concentrations of two sequential substrates (or products) in the pathway. Based on equations (4), (5) and (6), we deduce:

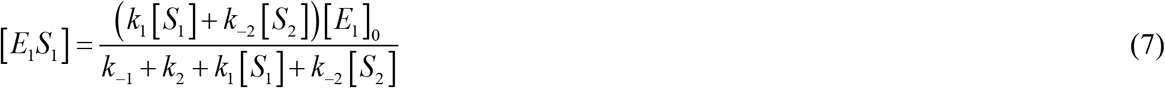

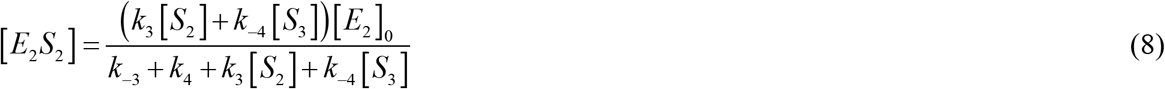

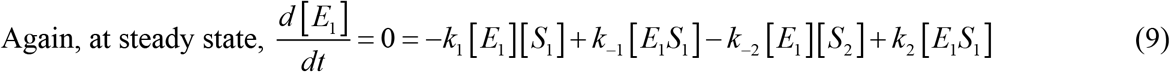

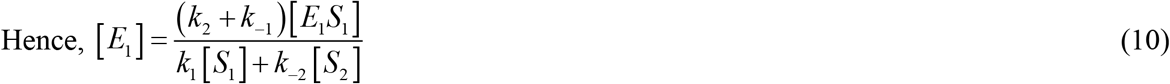

Similarly, one can show that:

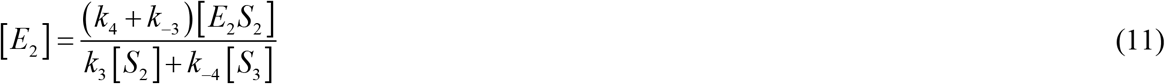

Since the pathway is at steady state, concentrations of reactants and products of every reaction remain unchanged with time, hence

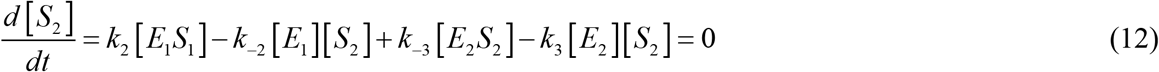

Using the expressions of [*E*_1_], [*E*_2_], [*E*_1_*S*_1_] and [*E*_2_*S*_2_] from equations (7), (8), (10) and (11) into equation (12),

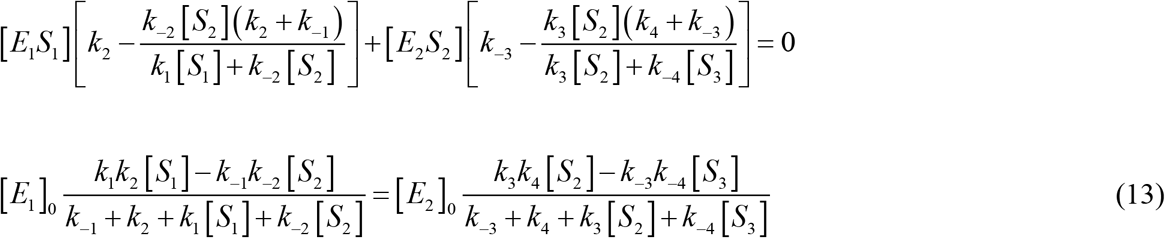

Using 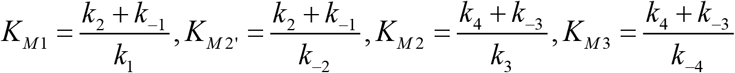, where *K*_*M* 1_ is the Michaelis constant of *E*_1_ for *S*_1_, *K*_*M* 2′_ is that of *E*_1_ for *S*_2_, *K*_*M* 2_ is that of *E*_2_ for *S*_2_ and *K*_*M* 3_ is that of *E*_2_ for *S*_3_, equation (13) can be written as:

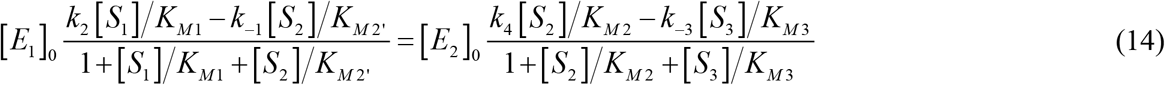

Assuming that *K*_*M* 2′_, *K*_*M* 3_ ≫ *K*_*M* 1_,*K*_*M* 2_ (in other words if *k*_−2_ and *k*_−4_ are very small) or the products have very low affinity back towards the enzyme, equation (14) reduces to the following:

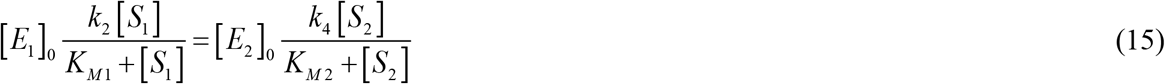

Equation (15) can be re-arranged to get the following hyperbolic or Michaelis-Menten like dependence of *S*_2_ on *S*_1_:

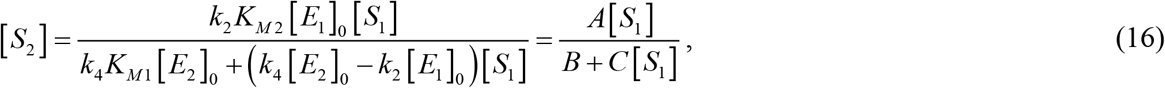

##### Case II: Sequential enzymes in pathway with Hill-like kinetics

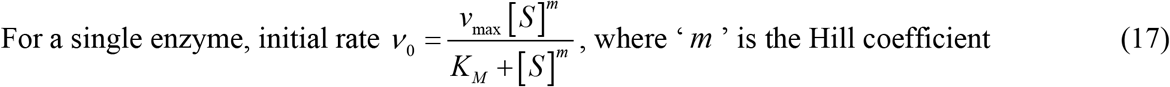

Now again consider the following scheme of sequential enzymes:

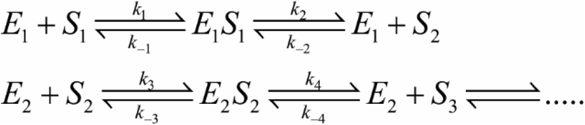

For reactants and products to be at steady state, earlier we derive Equation(15), which essentially equates the rate of production and consumption of *S*_2_ through the two enzymes (with the assumption that *k*_−2_ and *k*_−4_ are very small). In such a situation if either *S*_1_ or both *S*_1_ and *S*_2_ have limited diffusion, equation (15) can be written as the following based on equation (17):

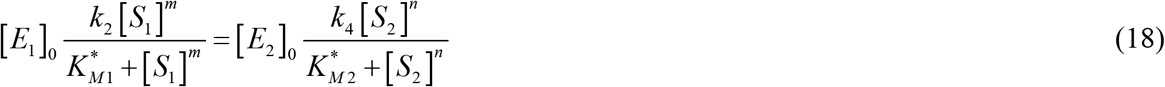

Where m and n are the Hill coefficient analogs of the two consecutive enzymatic steps.

Equation (18) can be rearranged as:

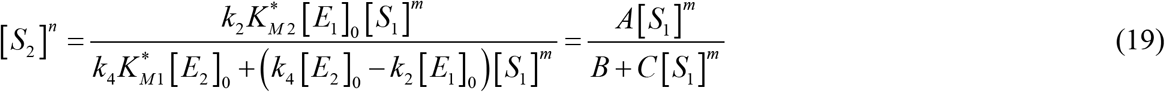

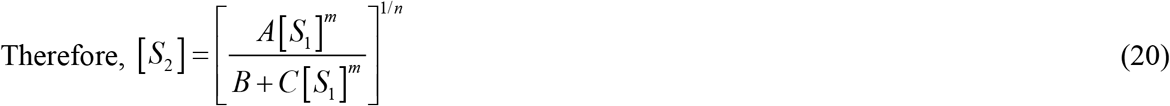

A numerical solution of equation (20) shows that *S*_2_ shows positive cooperativity as a function of *S*_1_ only if *m* > *n* (See Figure EV6).

#### Power law formalism for fractal kinetics

For a system where reactants and products diffuse freely, the rate constant of the reaction is time independent. However, under conditions of diffusion limitation (fractal kinetics), rate constant is no longer a constant, but varies with time in the following way:

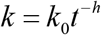

Where *h* is related to the fractal dimension of the medium.

In the following section, we adopt the power law formalism as shown in (Savageau, 1995) to convert the time dependent rate constant to a time independent one.

Considering the following simple reaction of two molecules of A forming a homodimer under conditions of diffusion limitation:

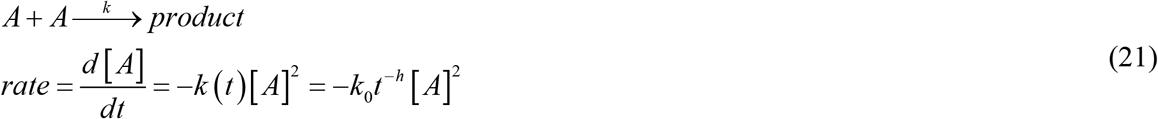

Integrating the above equation, we get [A] as a function of time

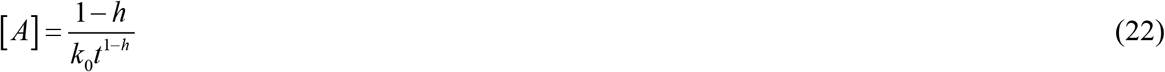

Rearranging this, we get

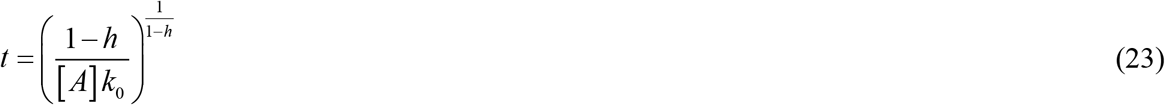

In the next step, we replace *t* in equation (21) with (23) to get:

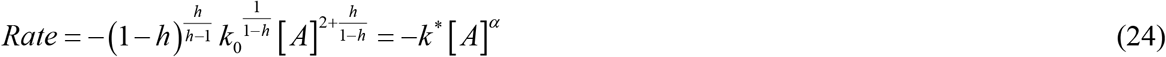

Where 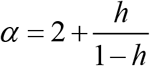 (Note that though this is a bimolecular reaction, the actual molecularity is >2 under fractal conditions)

#### Application for enzyme kinetics

Assume the following simple case:

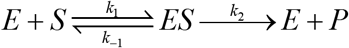

Consider that the substrate S is diffusion limited, hence *k*_1_ (a bimolecular rate constant) will be time dependent.

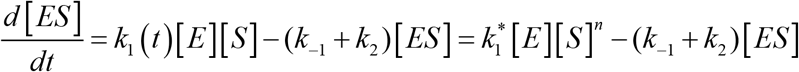

Where 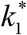 is the apparent time independent rate constant, and *n* is related to the fractal dimension of the medium.

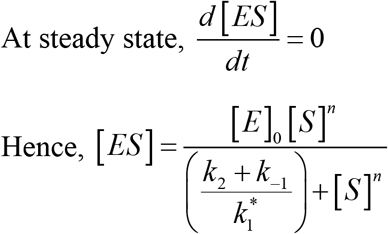

Rate of the reaction

Where 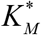 is the apparent Michaelis constant, and *n* is the Hill coefficient analog.

## Data availability

The datasets produced in this study are available in the following database: Metabolomics data: Metabolights MTBLS2795 (www.ebi.ac.uk/metabolights/MTBLS2795). The processed data are included in source data for Figures 2 and 5.

## Expanded View

The manuscript contains 6 Extended View figures (Figures EV 1–6).

### Expanded View Figure Captions

**Figure EV1:**
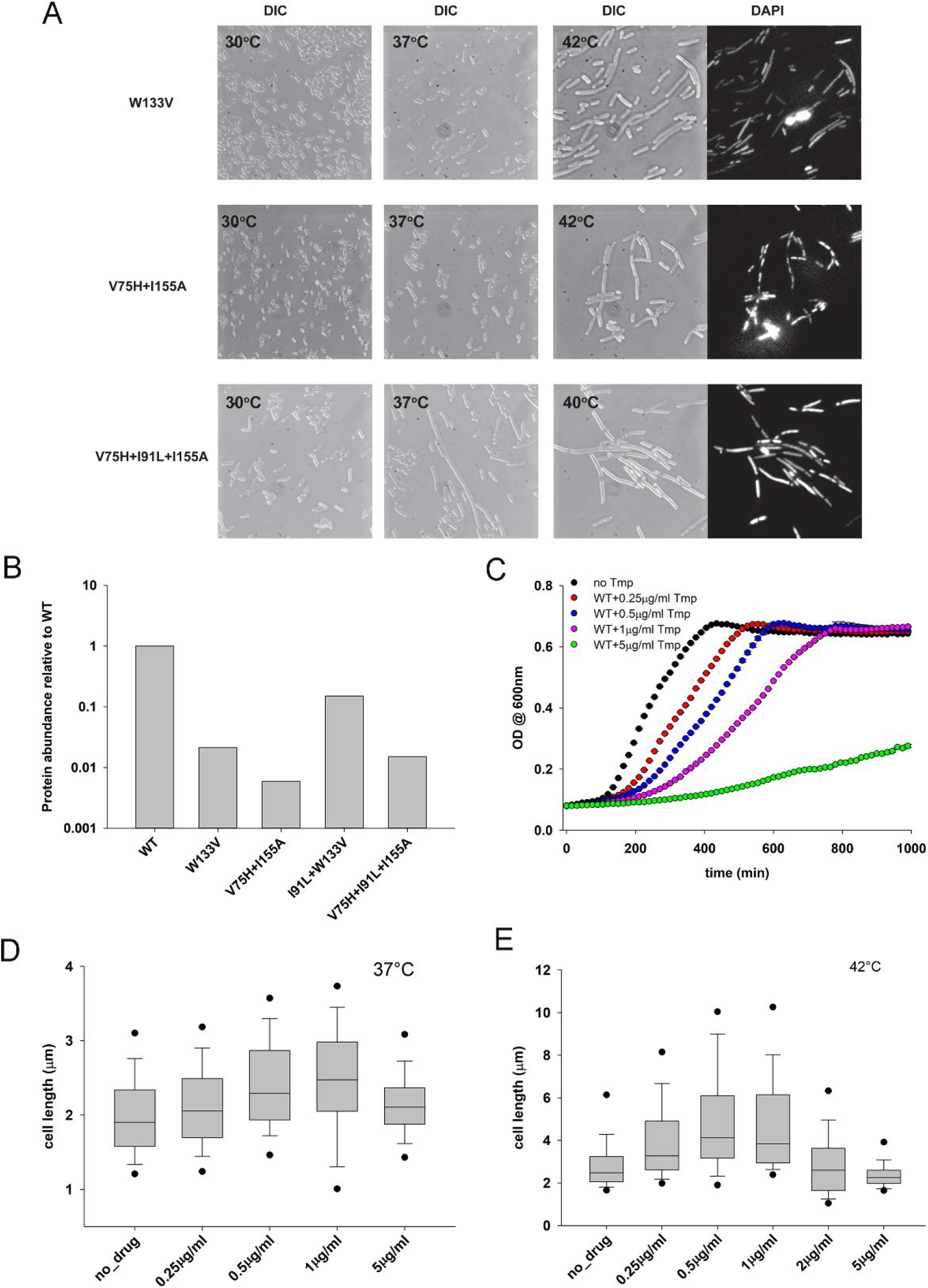
Filamentation of mutant DHFR strains and effect of Trimethoprim on filamentation. **(A)** Destabilizing mutations in DHFR induce filamentous phenotype. Live cells DIC images with DAPI nucleoids staining of W133V, V75H+I155A, and V75H+I91L+I155A DHFR *E. coli* MG1655 strains. Prior to microscopy, cells were grown at 30°C, 37°C, and 42°C (V75H+I91L+I155A was grown at 40°C) in amino acid supplemented M9 medium for 4 hours (see *Methods*). (B) Intracellular abundance of WT and mutant DHFRs measured by Western blot. WT, W133V and V75H+I155A were grown for 4 hours at 42°C while I91L+W133V and V75H+I91L+I155A strains were grown for 4 hours at 40°C in amino acid supplemented M9 medium before being harvested. The data is also reported in (Bershtein *et al*., 2015a). (C) The effect of WT DHFR inhibition by trimethoprim (Tmp) on growth. WT DHFR cells were grown at 42°C in amino acid M9 medium, and their growth was monitored by OD at 600nm. The data were fit to a 4-parameter Gompertz equation as described in (Bhattacharyya *et al*., 2017) to derive growth parameters. (D, E) Distribution of cell length of WT *E. coli* as a function of Tmp concentration when grown in amino acid supplemented M9 medium at (D) 37°C and (E) 42°C. Concentrations of Tmp slightly below or near the MIC (1μg/ml) results in maximum filamentation, while the effect dies down at higher concentrations. Filamentation is much more pronounced at 42°C than at 37°C. The central band in the box plots represents the median of the distribution, the box ends represent the 25^th^ and 75^th^ percentile, the whiskers represent the 10^th^ and 90^th^ percentile, while the dots represent the 5^th^ and 95^th^ percentile. Data was usually obtained from 2-3 biological replicates. The number of cells used to derive the boxplot distributions in the different panels range usually between 200-500.

**Figure EV2:**
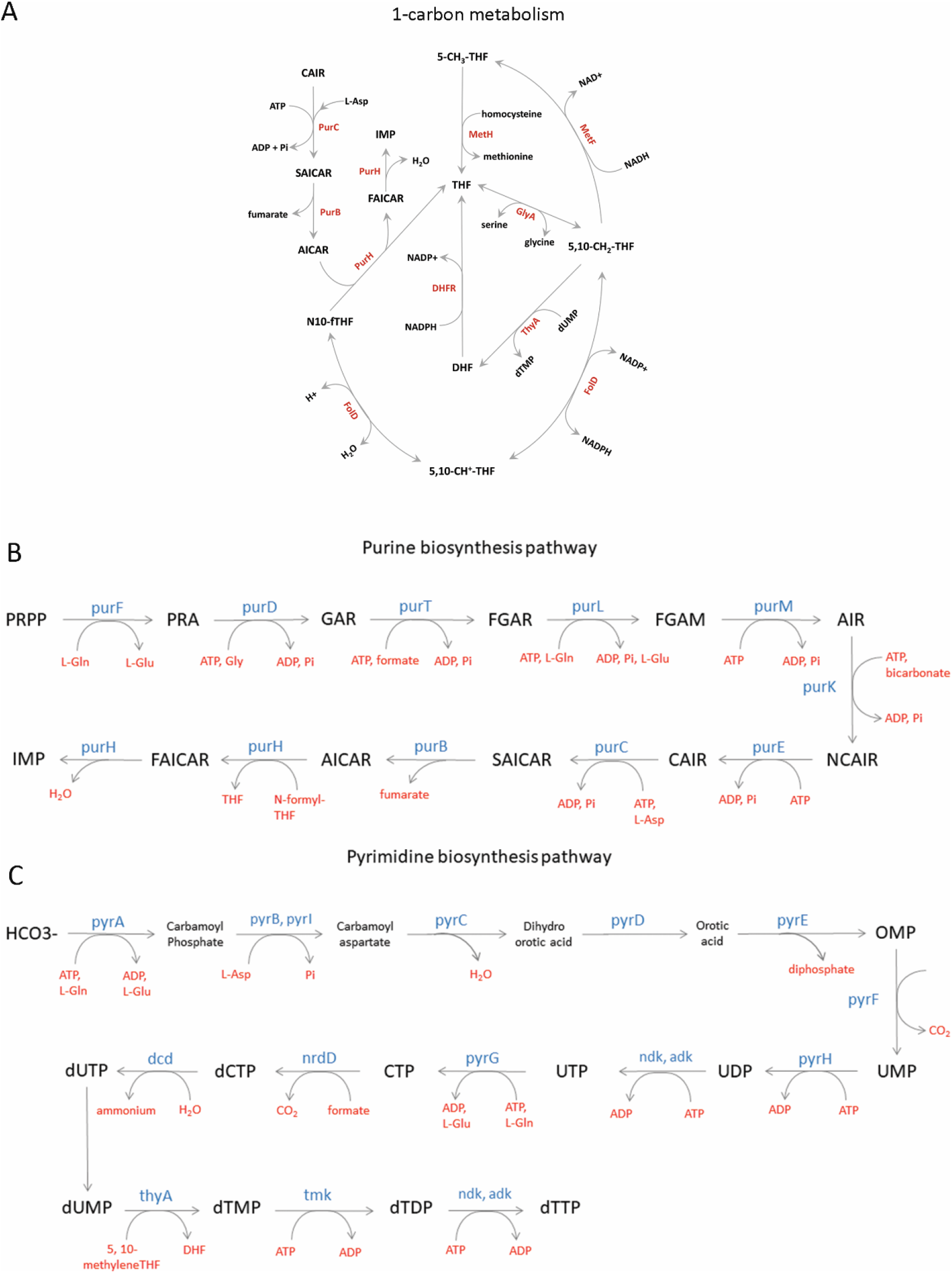
Pathways that use up product of DHFR activity. Schematic representation of (A) 1-carbon metabolism metabolism pathway (adapted from (Bhattacharyya *et al*., 2016)) (B) *de novo* purine biosynthesis pathway and (C) *de novo* pyrimidine biosynthesis pathway.

**Figure EV3:**
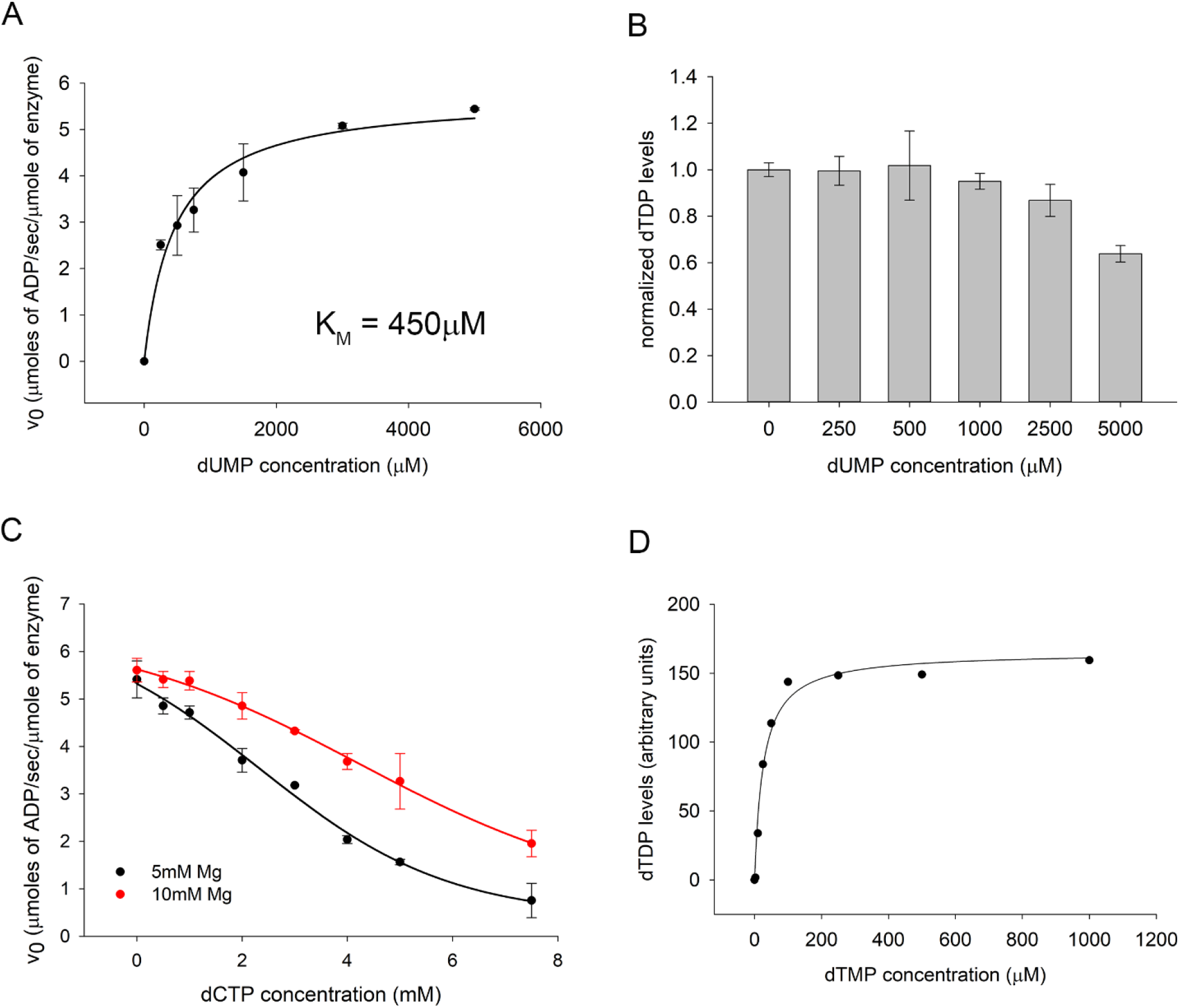
Effect of dUMP and dCTP on the *in vitro* activity assay of Thymidylate Kinase (Tmk). (A) Activity assay of purified Tmk enzyme using dUMP as substrate. ATP concentration is kept saturating at 1mM. The K_M_ for dUMP is 450μM, compared to 13μM for dTMP. (B) Activity assay of Tmk was carried out in the presence of 100mM dTMP and 1mM ATP, and varying concentration of the inhibitor dUMP. dTDP levels were measured by HPLC followed by mass spectrometry. The data was fitted with a 4-parameter sigmoid curve to obtain an apparent K_I_ of 3.9mM for dUMP. (C) Activity assay of Tmk was carried out in the presence of 100mM ATP and 1mM dTMP, and varying concentration of dCTP. ADP levels were measured using a NADH based coupled spectrophotometric assay. The red and black points indicate data acquired under different concentrations of Mg^2+^. The data were fitted with a 4-parameter sigmoid curve to obtain apparent K_I_ of 2.3mM and 4.2mM at 5 and 10mM Mg^2+^ concentrations respectively. For panels A to C, the error bars represent SD of three technical replicates. (D) Activity assay of purified Tmk as a function of dTMP concentration in the presence of 5mM dUMP and 2.5mM dCTP as inhibitors. ATP concentration was kept saturating at 1mM. The dTDP levels were monitored using HPLC followed by mass-spectrometry. Even in the presence of inhibitors, the activity data here conforms to MM kinetics.

**Figure EV4:**
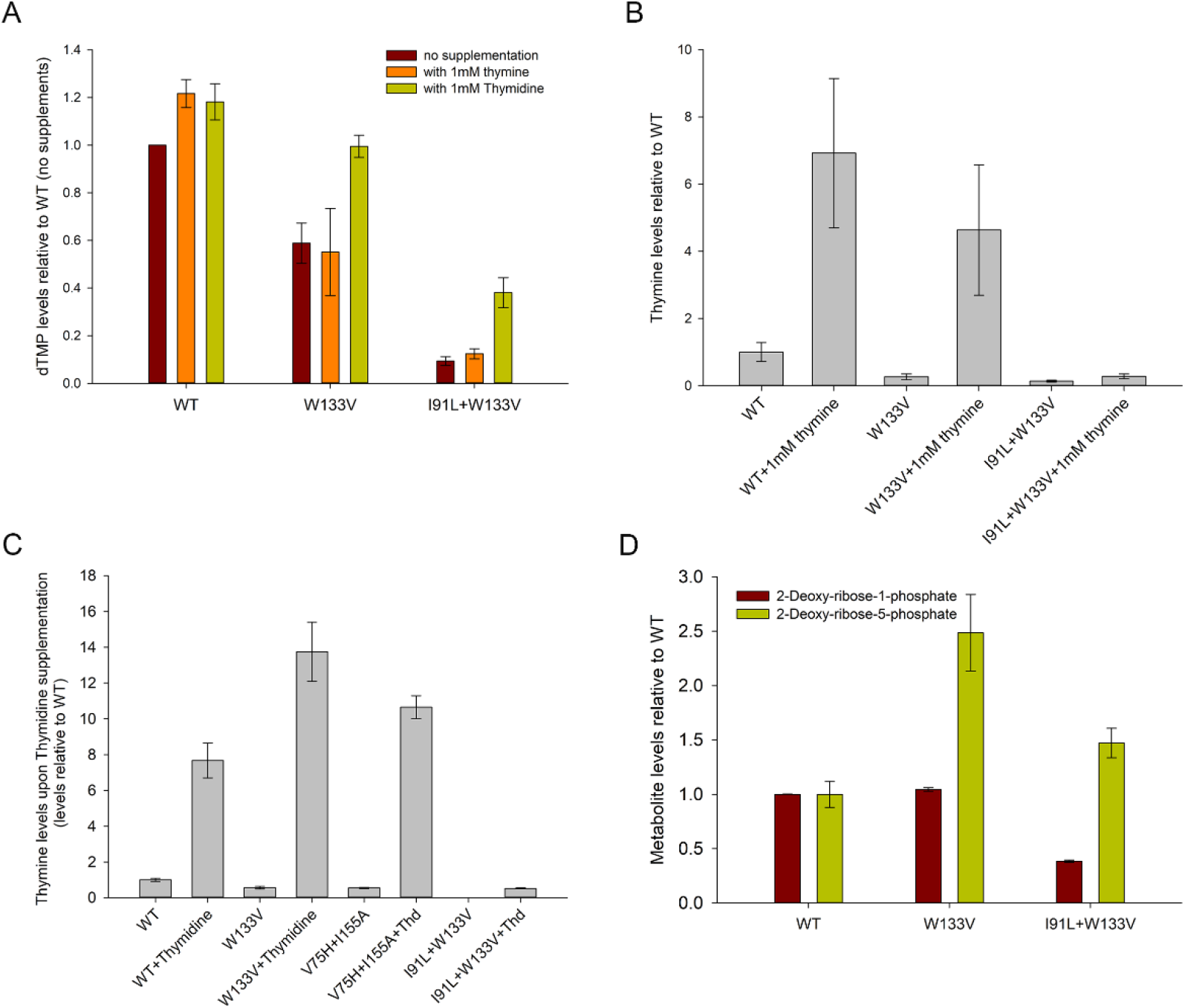
dTMP production through pyrimidine salvage pathway using thymidine and thymine supplementation. (A) Intracellular dTMP levels in WT and mutant strains upon addition of 1mM thymine or thymidine to the growth medium. Values are relative to those in WT strain (without any metabolite addition) after 4 hours of growth. Mutants show improvement in dTMP levels only upon thymidine addition. (B) Intracellular thymine levels in WT and mutant strains increase when grown in the presence of 1mM thymine in the medium, indicating that it is up taken by the cells. (C) Intracellular thymine levels in WT and mutant cells following growth with thymidine supplementation. Increase in thymine levels indicates substantial degradation of thymidine in the salvage pathway through DeoA enzyme. (D) Intracellular 2-deoxy-ribose-1-phosphate and 5-phosphate levels in WT and mutant cells. Mutants accumulate substantially high levels of the 5-phosphate variant, indicating its channeling into energy metabolism. For all panels, error bars represent SEM of at least three biological replicates.

**Figure EV5:**
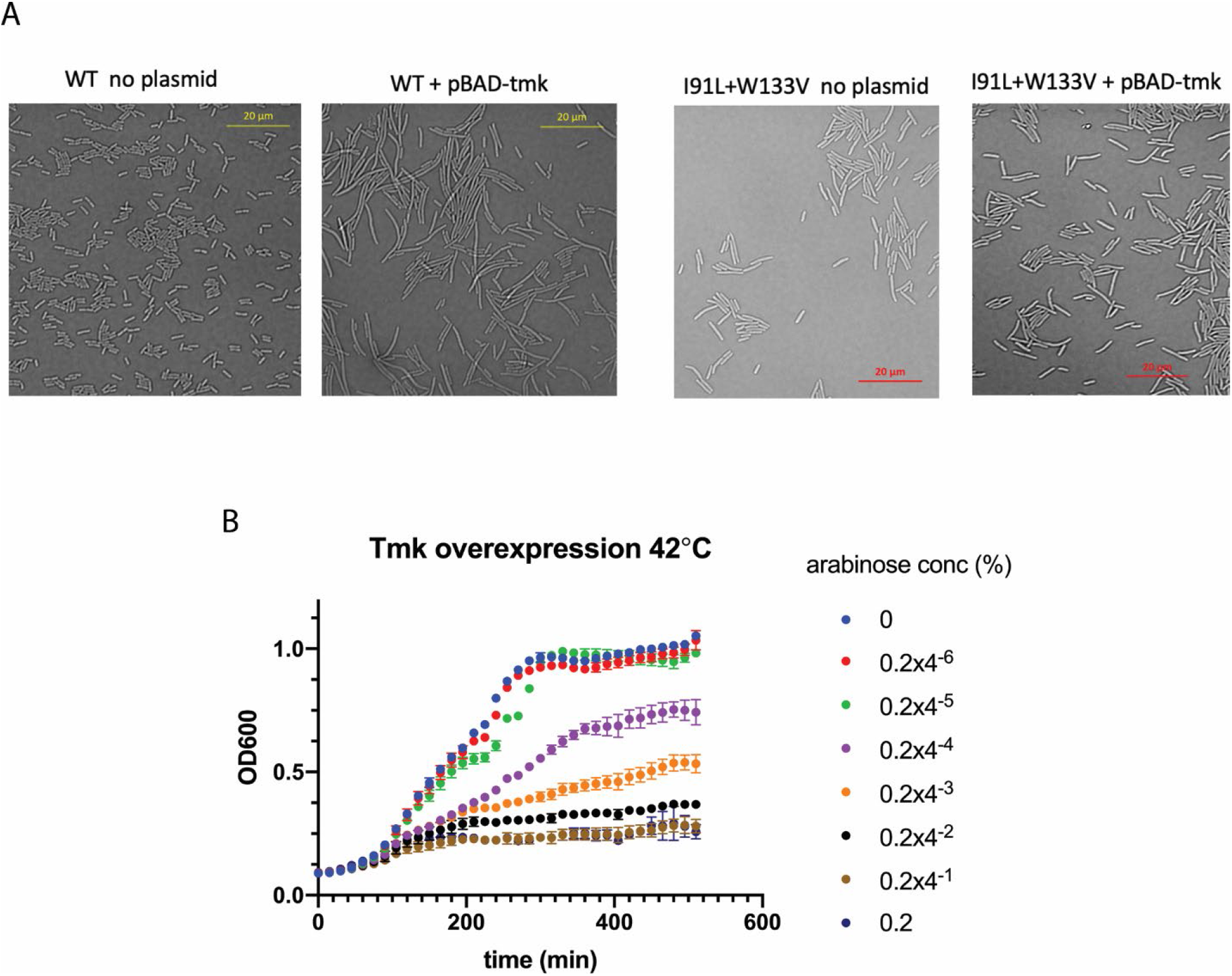
Effect of overexpression of Tmk on growth and morphology of WT and mutant DHFR strains. (A) DIC images of untransformed WT and I91L+W133V mutant cells as well as those transformed with pBAD plasmid that expresses Thymidylate Kinase under control of arabinose promoter. Cells were grown at 42°C for 4 hours (40°C for mutant) in amino acid supplemented M9 medium in the presence of 0.2% of arabinose. While expression of Tmk does not rescue filamentation of mutant cells, it produces filamentation of WT cells. (B) Growth curves of WT *E. coli* cells (BW27783) at 42°C following overexpression of Tmk from a pBAD plasmid with different concentrations of arabinose inducer. The data shows that overexpression of Tmk is toxic. Error bars represent SD of three technical replicates.

**Figure EV6:**
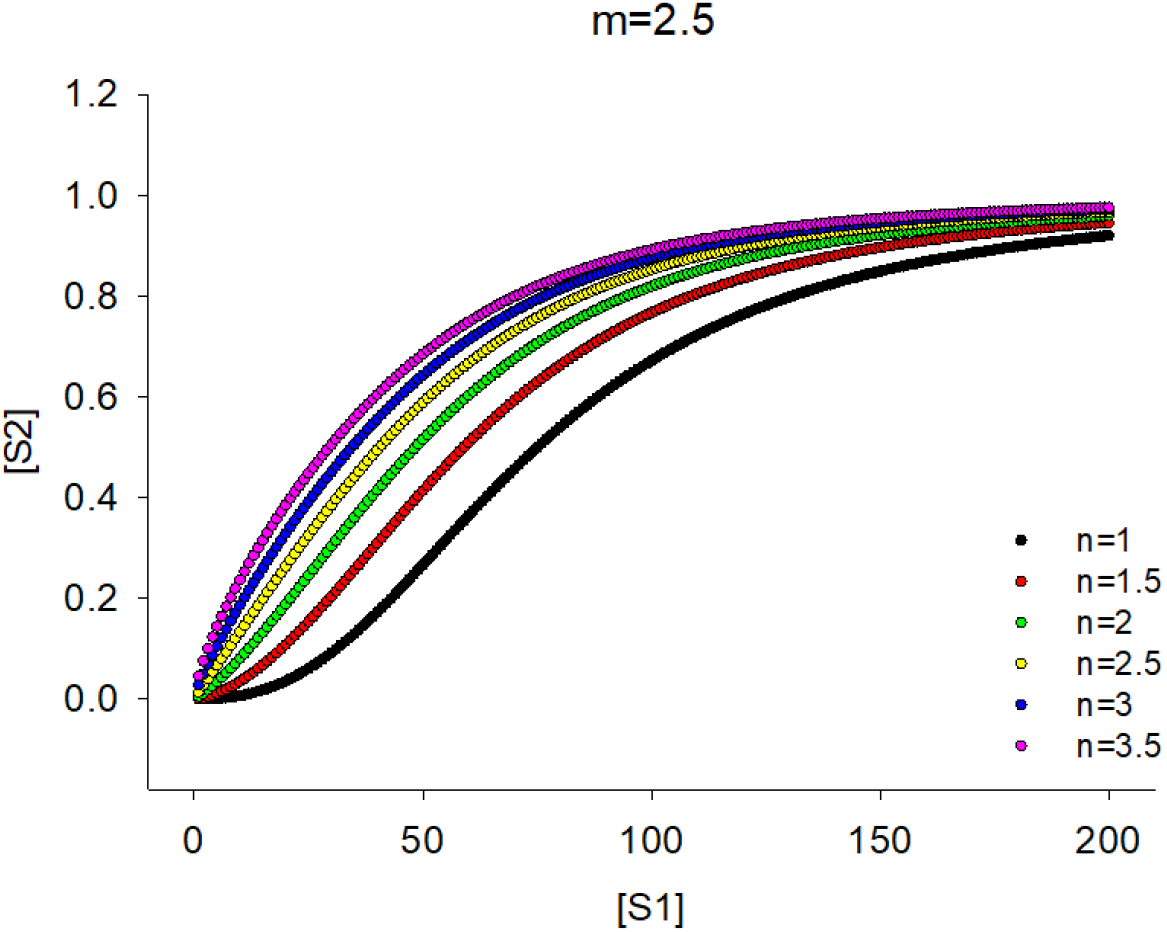
Numerical plot of Eq.20 showing steady state concentrations of a product and substrate in an enzymatic chain where sequential enzymes have Hill coefficients m=2.5 and variable n

## References

Adkar BV, Bhattacharyya S, Gilson AI, Zhang W, Shakhnovich EI (2019) Substrate inhibition imposes fitness penalty at high protein stability. Proc Natl Acad Sci U S A 116: 11265–11274

Adkar BV, Manhart M, Bhattacharyya S, Tian J, Musharbash M, Shakhnovich EI (2017) Optimization of lag phase shapes the evolution of a bacterial enzyme. Nat Ecol Evol 1: 149

Ahmad SI, Kirk SH, Eisenstark A (1998) Thymine metabolism and thymineless death in prokaryotes and eukaryotes. Annual review of microbiology 52: 591–625

An S, Kumar R, Sheets ED, Benkovic SJ (2008) Reversible compartmentalization of de novo purine biosynthetic complexes in living cells. Science 320: 103–106

Bazill GW (1967) Lethal unbalanced growth in bacteria. Nature 216: 346–349

Bennett BD, Kimball EH, Gao M, Osterhout R, Van Dien SJ, Rabinowitz JD (2009) Absolute metabolite concentrations and implied enzyme active site occupancy in Escherichia coli. Nat Chem Biol 5: 593–599

Bershtein S, Choi JM, Bhattacharyya S, Budnik B, Shakhnovich E (2015a) Systems-level response to point mutations in a core metabolic enzyme modulates genotype-phenotype relationship. Cell Rep 11: 645–656

Bershtein S, Mu W, Serohijos AW, Zhou J, Shakhnovich EI (2013) Protein Quality Control Acts on Folding Intermediates to Shape the Effects of Mutations on Organismal Fitness. Mol Cell 49: 133–144

Bershtein S, Mu W, Shakhnovich EI (2012) Soluble oligomerization provides a beneficial fitness effect on destabilizing mutations. Proc Natl Acad Sci U S A 109: 4857–4862

Bershtein S, Serohijos AW, Bhattacharyya S, Manhart M, Choi JM, Mu W, Zhou J, Shakhnovich EI (2015b) Protein Homeostasis Imposes a Barrier on Functional Integration of Horizontally Transferred Genes in Bacteria. PLoS Genet 11: e1005612

Bershtein S, Serohijos AW, Shakhnovich EI (2017) Bridging the physical scales in evolutionary biology: from protein sequence space to fitness of organisms and populations. Curr Opin Struct Biol 42: 31–40

Bhattacharyya S, Bershtein S, Shakhnovich EI (2017) Gene Dosage Experiments in Enterobacteriaceae Using Arabinose-regulated Promoters. Bio-Protocol 7: e2396

Bhattacharyya S, Bershtein S, Yan J, Argun T, Gilson AI, Trauger SA, Shakhnovich EI (2016) Transient protein-protein interactions perturb E. coli metabolome and cause gene dosage toxicity. Elife 5

Bhattacharyya S, Jacobs WM, Adkar BV, Yan J, Zhang W, Shakhnovich EI (2018) Accessibility of the Shine-Dalgarno Sequence Dictates N-Terminal Codon Bias in E. coli. Mol Cell 70: 894–905 e895

Davidi D, Noor E, Liebermeister W, Bar-Even A, Flamholz A, Tummler K, Barenholz U, Goldenfeld M, Shlomi T, Milo R (2016) Global characterization of in vivo enzyme catalytic rates and their correspondence to in vitro kcat measurements. Proc Natl Acad Sci U S A 113: 3401–3406

Dean AM, Thornton JW (2007) Mechanistic approaches to the study of evolution: the functional synthesis. Nat Rev Genet 8: 675–688

Deng Y, Gam J, French JB, Zhao H, An S, Benkovic SJ (2012) Mapping protein-protein proximity in the purinosome. J Biol Chem 287: 36201–36207

Dykhuizen DE, Dean AM, Hartl DL (1987) Metabolic flux and fitness. Genetics 115: 25–31

Fiehn O (2002) Metabolomics--the link between genotypes and phenotypes. Plant Mol Biol 48: 155–171

Frank SA (2013) Input-output relations in biological systems: measurement, information and the Hill equation. Biol Direct 8: 31

French JB, Jones SA, Deng H, Pedley AM, Kim D, Chan CY, Hu H, Pugh RJ, Zhao H, Zhang Y et al (2016) Spatial colocalization and functional link of purinosomes with mitochondria. Science 351: 733–737

Fuhrer T, Zampieri M, Sevin DC, Sauer U, Zamboni N (2017) Genomewide landscape of gene-metabolome associations in Escherichia coli. Mol Syst Biol 13: 907

Garcia-Contreras R, Vos P, Westerhoff HV, Boogerd FC (2012) Why in vivo may not equal in vitro - new effectors revealed by measurement of enzymatic activities under the same in vivo-like assay conditions. FEBS J 279: 4145–4159

Guenther S, Grobbel M, Lubke-Becker A, Goedecke A, Friedrich ND, Wieler LH, Ewers C (2010) Antimicrobial resistance profiles of Escherichia coli from common European wild bird species. Vet Microbiol 144: 219–225

Handakumbura PP, Stanfill B, Rivas-Ubach A, Fortin D, Vogel JP, Jansson C (2019) Metabotyping as a Stopover in Genome-to-Phenome Mapping. Sci Rep 9: 1858

Harrison BR, Wang L, Gajda E, Hoffman EV, Chung BY, Pletcher SD, Raftery D, Promislow DEL (2020) The metabolome as a link in the genotype-phenotype map for peroxide resistance in the fruit fly, Drosophila melanogaster. BMC Genomics 21: 341

Johnson CH, Ivanisevic J, Siuzdak G (2016) Metabolomics: beyond biomarkers and towards mechanisms. Nat Rev Mol Cell Biol 17: 451–459

Kitagawa M, Ara T, Arifuzzaman M, Ioka-Nakamichi T, Inamoto E, Toyonaga H, Mori H (2005) Complete set of ORF clones of Escherichia coli ASKA library (a complete set of E. coli K-12 ORF archive): unique resources for biological research. DNA Res 12: 291–299

Kopelman R (1988) Fractal reaction kinetics. Science 241: 1620–1626

Kwon YK, Higgins MB, Rabinowitz JD (2010) Antifolate-induced depletion of intracellular glycine and purines inhibits thymineless death in E. coli. ACS Chem Biol 5: 787–795

Kwon YK, Lu W, Melamud E, Khanam N, Bognar A, Rabinowitz JD (2008) A domino effect in antifolate drug action in Escherichia coli. Nat Chem Biol 4: 602–608

Li HQ, Chen SH, Zhao HM (1990) Fractal mechanisms for the allosteric effects of proteins and enzymes. Biophys J 58: 1313–1320

Liebovitch LS, Fischbarg J, Koniarek JP, Todorova I, Wang M (1987) Fractal model of ion-channel kinetics. Biochim Biophys Acta 896: 173–180

Lunzer M, Miller SP, Felsheim R, Dean AM (2005) The biochemical architecture of an ancient adaptive landscape. Science 310: 499–501

Maguin E, Brody H, Hill CW, D’Ari R (1986) SOS-associated division inhibition gene sfiC is part of excisable element e14 in Escherichia coli. J Bacteriol 168: 464–466

Møllgaard H, Neuhard J (1983) Biosynthesis of deoxythymidine triphosphate. Academic Press, New York, NY

Mulleder M, Calvani E, Alam MT, Wang RK, Eckerstorfer F, Zelezniak A, Ralser M (2016) Functional Metabolomics Describes the Yeast Biosynthetic Regulome. Cell 167: 553–565 e512

Nelson DJ, Carter CE (1969) Purification and characterization of Thymidine 5-monophosphate kinase from Escherichia coli B. J Biol Chem 244: 5254–5262

Patti GJ, Yanes O, Siuzdak G (2012) Innovation: Metabolomics: the apogee of the omics trilogy. Nat Rev Mol Cell Biol 13: 263–269

Pritchard RH, Zaritsky A (1970) Effect of thymine concentration on the replication velocity of DNA in a thymineless mutant of Escherichia coli. Nature 226: 126–131

Rodrigues JV, Bershtein S, Li A, Lozovsky ER, Hartl DL, Shakhnovich EI (2016) Biophysical principles predict fitness landscapes of drug resistance. Proc Natl Acad Sci U S A 113: E1470–1478

Rodrigues JV, Shakhnovich EI (2019) Adaptation to mutational inactivation of an essential gene converges to an accessible suboptimal fitness peak. Elife 8

Sangurdekar DP, Hamann BL, Smirnov D, Srienc F, Hanawalt PC, Khodursky AB (2010) Thymineless death is associated with loss of essential genetic information from the replication origin. Mol Microbiol 75: 1455–1467

Sangurdekar DP, Zhang Z, Khodursky AB (2011) The association of DNA damage response and nucleotide level modulation with the antibacterial mechanism of the anti-folate drug trimethoprim. BMC Genomics 12: 583

Savageau MA (1995) Michaelis-Menten mechanism reconsidered: implications of fractal kinetics. J Theor Biol 176: 115–124

Savageau MA (1998) Development of fractal kinetic theory for enzyme-catalysed reactions and implications for the design of biochemical pathways. Biosystems 47: 9–36

Schnell S, Turner TE (2004) Reaction kinetics in intracellular environments with macromolecular crowding: simulations and rate laws. Prog Biophys Mol Biol 85: 235–260

Sliusarenko O, Heinritz J, Emonet T, Jacobs-Wagner C (2011) High-throughput, subpixel precision analysis of bacterial morphogenesis and intracellular spatio-temporal dynamics. Mol Microbiol 80: 612–627

Torre-Bueno JR (1976) Temperature regulation and heat dissipation during flight in birds. J Exp Biol 65: 471–482

van Eunen K, Kiewiet JA, Westerhoff HV, Bakker BM (2012) Testing biochemistry revisited: how in vivo metabolism can be understood from in vitro enzyme kinetics. PLoS Comput Biol 8: e1002483

van Eunen KB, B. M. (2014) The importance and challenges of in vivo-like enzyme kinetics. Perspectives in Science 1: 126–130

Zampieri M, Sauer U (2017) Metabolomics-driven understanding of genotype-phenotype relations in model organisms. Current Opinion in Systems Biology 6: 28–36

Zaritsky A, Pritchard RH (1973) Changes in cell size and shape associated with changes in the replication time of the chromosome of Escherichia coli. Journal of bacteriology 114: 824–837

Zaritsky A, Woldringh CL, Einav M, Alexeeva S (2006) Use of thymine limitation and thymine starvation to study bacterial physiology and cytology. J Bacteriol 188: 1667–1679

Zotter A, Bauerle F, Dey D, Kiss V, Schreiber G (2017) Quantifying enzyme activity in living cells. J Biol Chem 292: 15838–15848

